# Using extreme value statistics to reconceptualize psychopathology as extreme deviations from a normative reference model

**DOI:** 10.1101/2022.08.23.505049

**Authors:** Charlotte Fraza, Mariam Zabihi, Christian F. Beckmann, Andre F. Marquand

## Abstract

For many problems in neuroimaging, the most informative features occur in the tail of the distribution. For example, when considering psychiatric disorders as deviations from a ‘norm’, the tails of the distribution are considerably more informative than the bulk of the distribution for understanding risk, for stratifying and predicting such disorders and for anomaly detection. Yet, most statistical methods used in neuroimaging focus on modelling the bulk and fail to adequately capture extreme values occurring in the tails. To address this, we propose a framework that combines normative models with multivariate extreme value statistics to accurately model extreme deviations of a reference cohort for individual participants. Normative models are now widely used in clinical neuroscience and similar to the employment of normative growth charts in pediatric medicine to track a child’s weight in relation to their age, normative models can be used with neuroimaging measurements to quantify individual neurophenotypic deviations from a reference cohort. However, formal statistical treatment of how to model the extreme deviations from these models has been lacking until now. In this article we provide such an approach, inspired by applications of extreme value statistics in meteorology. Since the presentation of extreme value statistics is usually quite technical, we begin with a non-technical introduction to the fundamental principles of extreme value statistics, to accurately map the tails of the normative distribution for biological markers, including mapping multivariate tail dependence across multiple markers. Next, we give a demonstration of this approach to the UK Biobank dataset and demonstrate how extreme values can be used to accurately estimate risk and detect atypicality. This framework provides a valuable tool for the statistical modelling of extreme deviations in neurobiological data, which could provide us with more accurate and effective diagnostic tools for neurological and psychiatric disorders.

## Introduction

In many domains, understanding the extreme values of a given variable is of great importance since they have the most significant impact on outcomes of interest (1–3). In neuroscience, we are often similarly interested in the extreme values of a distribution. There are several reasons for this. First, many neurological and psychiatric disorders are characterized by patterns of deviations in specific regions or circuits of the brain (4–6). These deviations may be difficult to detect using traditional statistical methods that focus on the mean or median values of the data. However, they may be more readily apparent in the extreme values of the data, which reflect significant deviations from the norm. For example, we know that atypical brain development underlies many brain disorders (7,8). If we understand early deviations in brain development better, we could provide risk factors for developing several mental disorders. For example, early extreme deviations in frontal cortex development from the norm could be a risk factor for attention deficit disorder (9). Or, an extreme sudden decrease of dopamine cells in the substantia nigra could be an early biomarker for Parkinson’s disease (7). Small variations in brain development are generally not cause for concern, as they are within the normal range of individual differences. However, extreme deviations from normal brain development in multiple brain regions can be indicative of underlying neurological or psychiatric conditions that can affect a person’s ability to function in their daily life. By accurately mapping these extreme deviations, researchers can gain a better understanding of the brain’s structure and function and identify potential risk factors for psychiatric disorders such as depression, schizophrenia, and ADHD. Second, many neurological and psychiatric disorders are complex and heterogeneous, with multiple subtypes and comorbidities (10,11). By focusing on the tails of the distribution across multiple variables, researchers can identify patterns of tail dependence that might inform about distinct subtypes of the on the basis of brain activity or structure. Finally, the tails of the distribution might also provide particularly useful information for gauging prognosis and treatment response. For instance, people with more extreme values in particular brain regions may respond better to certain types of treatment or may have worse outcomes overall.

Reliable estimation of the tails of a distribution poses multiple challenges. Firstly, data in the tails of a distribution are, by definition, sparse, which makes it difficult to acquire a sufficient number of observations to draw meaningful conclusions (12). Secondly, classical approaches in statistics are designed to focus on the central part (i.e. bulk) of the distribution rather than the tails. This makes it difficult to capture the subtle nuances of the tail behavior, leading to inaccurate estimation of tail probabilities (13). This is crucial because for many applications underestimating or overestimating the likelihood of extreme events can lead to large errors in risk assessment and treatment decisions. Accurate estimation of tail probabilities is, therefore, critical for obtaining reliable estimates of risk and optimal clinical decision-making. An additional problem is that extreme values usually do not occur in isolation, therefore modelling dependencies in the tails of multiple variables is also important for accurate inference. In this respect, it is crucial to recognise that tail dependence does not in general correspond to correlation in the traditional sense. For instance, two variables may be highly correlated, yet show low tail dependence and vice versa. We illustrate this problem didactically using a watershed model inspired by hydrology in the next section.

In this paper, we propose a neuroimaging framework for understanding and exploring patterns in extreme brain deviations for the individual, based on extreme value statistics and normative modelling. Normative modelling provides statistical estimates of the distribution of a given brain measure across the population and can be used to estimate individual risk or deviation scores (8,14). Combining the strength of normative modelling with extreme value statistics will allow us to accurately estimate extreme deviation risk scores for the individual. The combination of these methods enables us to establish brain-behavior mappings that are more sensitive to the multivariate structure underlying the extreme deviations from an expected pattern (e.g., normative brain development). We predict that the accurate understanding of small probabilities of extreme events is crucial for understanding several types of behavior and psychiatric disorders.

The main goals of this paper are: first, to provide a didactic introduction to extreme value statistics along with guidelines for implementing them in practice, which is important because extreme value statistics is quite a technical topic. Second, we provide a proof of principle, to demonstrate the use of multivariate extreme value statistics for neuroimaging data. Whilst we anticipate that extreme value statistics is widely useful in neuroimaging, it is particularly naturally suited to normative modelling, because normative models provide a flexible method to map multiple variables that might have different non-Gaussian distributions (e.g. brain volume and cortical thickness) to z-statistics, which have the same asymptotically Gaussian distribution. This is particularly important for calibrating and fitting these models.

In summary, our main contributions are: (i) Developing a comprehensive analytical framework for multivariate extreme value statistics for neuroimaging data; (ii) applying and demonstrating it to the UK Biobank dataset; (iii) establishing clear underlying of tail pairwise dependence between several brain phenotypes; (iv) summarizing the tail pairwise dependence using principal components based on the extreme values, henceforth called extreme principal components. Finally, (v) we relate these extreme principal components to several behavioral phenotypes.

## 1. Theoretical background

### 1.1 Normative models

Normative modelling is a method for understanding and modelling the heterogeneity in the population. Similar to pediatric growth charts, which are used to model the distribution of height or weight of children according to their age and sex, normative models can be used to model neuroimaging data according to a set of clinical or demographic covariates (e.g. age, sex and site) (14,15). The outputs of this analysis are the median and centiles of variation of the data as a function of the covariates. Using normative models, we shift away from the case-control setting and are able to make inferences of deviation from a reference cohort for each individual. Normative models have been used extensively to create large reference cohorts, establish individual deviation scores from these reference cohorts and link the deviation scores to diverse psychiatric problems (4–6,16,17). We typically aim to use the deviation scores from various structural and functional brain measures as features that are then associated with clinical or behavioural scores. Further details about the normative modelling approach employed here are provided in the supplement.

### 1.2 Conceptualizing multivariate deviations using a watershed model

Most models of mental disorders conceptualize them as the emerging end product of aggregated causes, from disruptions in genetic pathways, brain networks, to the environment (18). For example, multiple neural system dysfunctions have been recorded that can lead to disruptions in attention, memory, and emotion regulation (19). Small individual disturbances to certain nodes in the brain network can cause combined effects that surpass a liability threshold for a specific or multitude of mental disorders. A complex psychiatric state such as schizophrenia, major depression or anxiety disorder is then thought to be caused by the combined heterogeneous deviations from typical network functioning or by one local disruption in the network. To illustrate this point, we give an analogy inspired by hydrology, where extreme value statistics are commonly used, see Figure 1. Under this analogy In order to understand the effects of a flooding event (e.g. at the river delta), predicting the frequency and intensity of extreme values at different nodes on the river becomes much more important than predicting the change in the mean, because the combined extreme events are what cause the most damage (20). We illustrate this in Figure 1, where we give a visual representation of the different types of extreme data that can exist at individual gauging stations that together cause a flood. It could be that one river in the network is having local water elevation, due to, for example, snow melting (Figure 1. A). This local water elevation then causes flooding downstream at the river basin. Or it could be that the water elevation is raised at multiple locations due to a more generalized effect, e.g. a storm across the land (Figure 1. B and C). Under this watershed model it becomes clear that it is important that we model the extremal dependence structure between the different gauging stations and water flow correctly, as it is not only the local extreme events, but also a co-occurrence of multivariate extremes that determines how severe the flooding will be (21,22).

**Figure 1.**
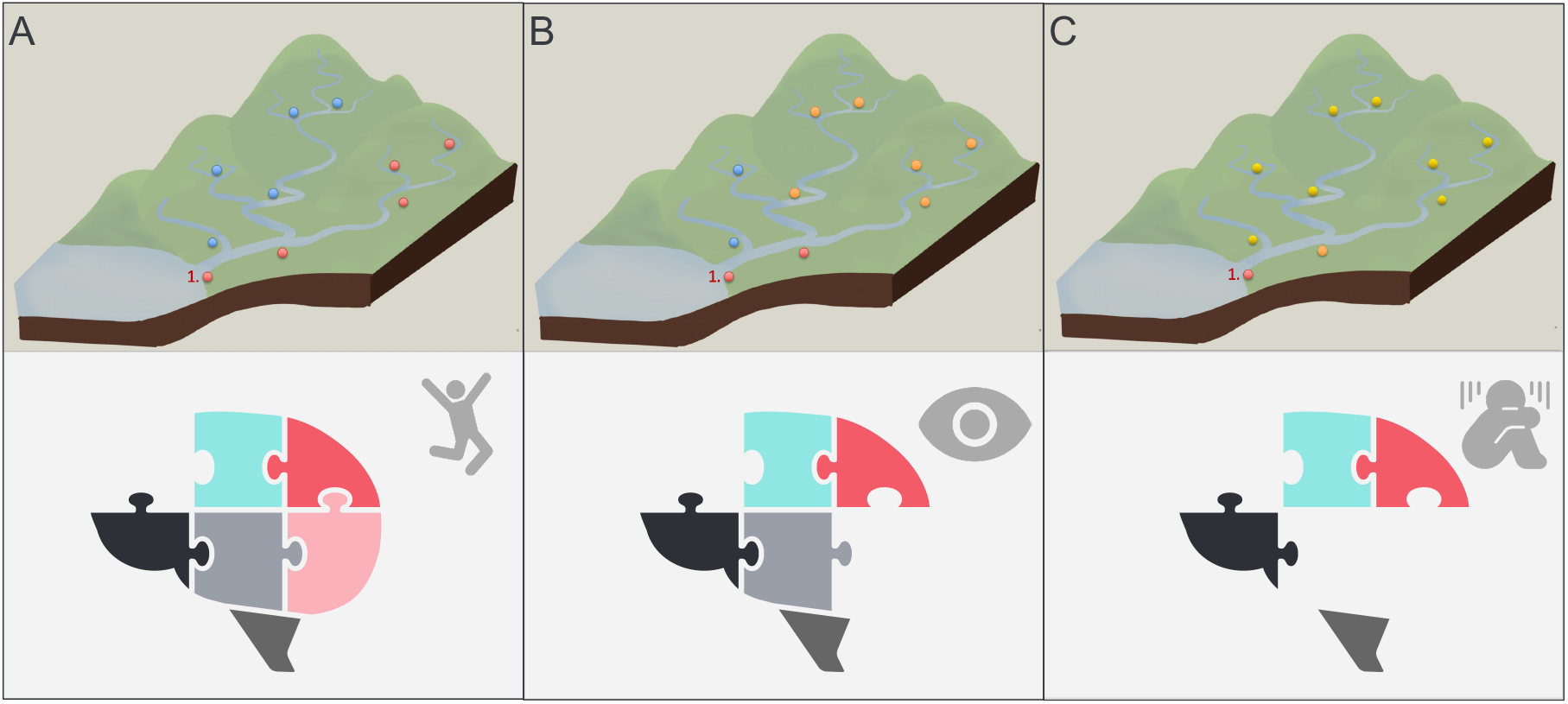
Visual example of different types of extreme river elevations that can cause local flooding at location 1. In panel A local elevation along one water bank gives rise to the flooding of the river basin at location 1. In panels B and C, it is the multivariate extreme elevation of the rivers in combination that gives rise to the flooding at location 1. Likewise, extremes in behavior can be understood as being the end product of several large deviations in certain brain regions or networks.

This analogy can be transferred to psychopathology, as proposed in Cannon’s watershed model (18). Under this model, pathophysiological brain states can manifest either due to local brain disruptions in certain brain regions or nodes in the network or via a more widespread disruption in connected brain systems. The widespread disruption can cause psychopathology by a ‘downstream’ aggregation of different effects. Thus, there are many different ways that small causal effects (the tributaries) can be aggregated to yield the full-blown expression of a disorder (the river delta). The key points are that: (i) neural correlates of psychopathology can be understood as manifesting at the extreme end of continuous axes of variation across the population, (ii) these extreme deviations may be spread across multiple brain systems and may not necessarily overlap across individuals and (iii) these deviations may not be reflected in only one data modality. We now show how this model can be operationalized within the context of multivariate extreme value statistics.

### 1.3 Introduction to extreme value statistics

We will rely on extreme value statistics to accurately model probabilities of the tails of the distribution (23), that is, extreme events. We start with exploring a univariate extreme value model fitting method which serves as a building block for the multivariate extreme value statistics approach we introduce later. There are two main motivations for this. First, depending on the question of the user the univariate approach could be sufficient. For example, if one suspects that the extremes in *one* brain region are most predictive of behavior, such as the amygdala in generalized anxiety disorder (24). Second, threshold selection for multivariate extreme value statistics is most easily done by looking at individual model fit in univariate space, as we outline below.

For the multivariate extreme values, this work employs a framework that is specifically created for exploring extreme tail dependence when the dimensions become large, such as when we look at multiple brain regions or voxels simultaneously. We employ the tail pairwise dependence matrix (TPDM) (25) to understand the dependence structure in the extremes, which is similar to correlation or covariance variance matrix, but specified for the tails. We then compute an eigendecomposition of this matrix, similar to what is done in principal components analysis (PCA). This is motivated by the fact that, whilst standard covariance PCA estimates variation about the mean, it is not well-suited to modelling multivariate tail dependency for the extremes, because dependence in the bulk of a multivariate distribution may not be equivalent to dependence in the tails (25). The TPDM is similar to a covariance matrix, but instead of describing the dependence structure around the mean of the distribution, the TPDM captures the dependence structure in the joint tails. An eigendecomposition gives us an ordered basis of eigenvectors based on the extremes, and values that capture, in order, the most important vectors for explaining the extreme dependence structure (25,26). We call the method ‘TPDM PCA’ or ‘extreme PCA’ and the associated principal components (PCs) ‘TPDM PCs’ or ‘extreme PCs’. Similar to standard PCs the TPDM PCs can be interpreted by looking at the individual contributions of each variable to the TPDM PCs, for example, certain regions of interest (ROIs) in neuroimaging data. This gives an indication of which variables are most important in understanding the extreme deviations.

## 2. Materials and methods

### 2.1 Data Sample

The data used in the paper came from the UK Biobank imaging data set (28,29). We used the image-derived phenotypes (IDPs), which are pre-processed imaging data, generated by the image-processing pipeline developed and run on behalf of the UK Biobank (30). These are summary measures of several imaging modalities, including structural, functional and diffusion modalities, making them ideally suited for evaluating new methods and theories. For the full details on the pre-processing steps and the design of the study, the following document can be read ‘UK Biobank Brain Imaging Documentation’ (31). We used data from 43,893 participants between 40 and 80 years old, with around 53% females. For the exact distribution of the age range and the number of participants currently in our dataset, see Supplement 1. For this study, we pre-processed the IDPs using the FUNPACK software (32). We used 2006 IDPs in total, including structural, functional and diffusion MRI IDPs processed by the UKB consortium. We split the available IDP data into a training set and a test set with a 10-90 split, giving us 4405 participants for training and 39488 participants for testing. This was a deliberate choice as we aimed to retain a large number of testing subjects to evaluate the new multivariate extreme value framework on. For the UK Biobank, the dataset is very large, thus the number of training subjects is still acceptable; however, with a smaller dataset, we would recommend a more conservative split (e.g. split-half).

For the behavioural analysis, the UK Biobank dataset provided an extensive number of behavioral, clinical, lifestyle and cognitive variables (28). These non-imaging derived phenotypes (nIDPs) are categorized within FUNPACK into seven different categories: cognitive phenotypes (199 variables), lifestyle environmental alcohol (9 variables), lifestyle environment tobacco (4 variables), lifestyle environment general (44 variables), lifestyle environment exercise work (37 variables), mental health (29 variables), neurodegenerative disorders (6 variables). For simplicity we follow this categorization here.

### 2.2 Overview of analytic workflow

In Figure 2, we present a high-level overview of all the steps in the multivariate extreme value framework we propose. In brief, we first estimated a normative model for all the IDPs using age, sex, and site as our covariates. We then applied a warped Bayesian Linear Regression (BLR) model, which helped us to predict the neuroimaging data from the covariates, whilst modelling non-Gaussianity and providing us with posterior predictive intervals for each prediction. This essentially follows an approach we have validated previously (6,33). We then evaluated the performance of the BLR model and the likelihood warping by examining multiple error metrics, namely explained variance, mean standardized log loss (MSLL), kurtosis and skewness (see Supplement for details). Then, we collected the deviation or *z*-scores for each participant and all the IDPs into one *z*-scores matrix including all the IDPs we modeled. Next, we transformed the marginals and converted to polar coordinates, which are standard operations in extreme value statistics and will be explained in more detail below. Then, we thresholded the data at the 95-quantile in polar coordinates. Finally, the TPDM was calculated, based on the extreme quantiles of the data. On this matrix, we performed an eigenvalue decomposition, which gave us an eigenbasis that explains most of the extremal structure in the first few eigenvectors. Finally, we correlated the first few extreme PCs back to behavior, to establish a brain-behavior relationship. These steps are explained in more detail in the next sections.

**Figure 2.**
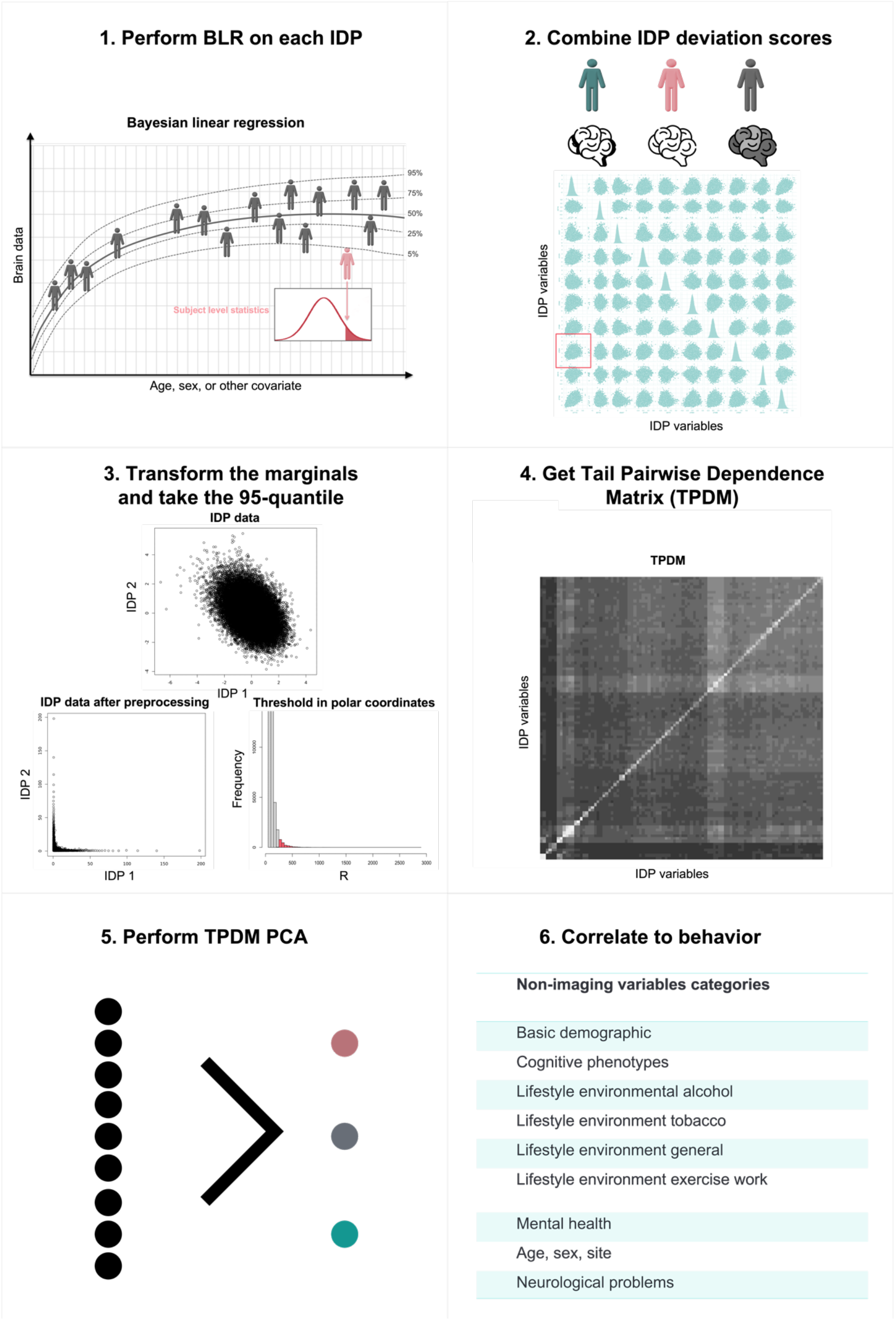
Overview of the proposed extreme normative modelling approach showing all the steps in the pipeline. 1. Estimate a normative model with likelihood-warped Bayesian Linear regression for each (neuro) image-derived phenotype (IDP). This allows us to create deviation or z-scores from the mean trend for each participant. 2. Combine the z-scores of all the participants and each IDP into one matrix. 3. Transform the marginals of the data such that the marginal distribution is varying with tail index = 2 and threshold the data in polar coordinates at the 95-quantile. 4. Calculate the tail pairwise dependence matrix (TPDM) based on the outer 95-quantile of the z-scores. This matrix captures the extreme structure in the outer tails. 5. Perform an eigenvalue decomposition on the TPDM, creating an independent basis on which the first few eigenvectors contain most of the information of the extremal structure. 6. Correlate the first few extreme principal components to behavior to indicate how multivariate extreme values influence behavior. See the text for full details.

### 2.4 Extreme value statistics

If we consider the entire distribution of deviation scores, we can approximate it as being composed of two components: the bulk of the distribution and the tails. Usually, we can fit a parametric model to the bulk of the distribution, such as a Gaussian distribution. However, if we are interested in the far upper and lower tails or the extreme quantiles of a distribution, we have to apply extreme value statistics, which allows us to model the tail probabilities accurately (23,27). The extreme events in the tails have a small probability to occur, but usually exercise a large influence on relevant outcomes. However, the small occurrence or sparseness of measurements in the tails makes it difficult to model accurately. In the univariate case, two methods can be employed to address this, i.e. the ‘block maxima’ or the ‘exceedance’ (a.k.a. ‘peaks over threshold’) approach. We focus on the ‘peaks over threshold’ approach in this paper, which is summarized in Figure 3.

**Figure 3.**
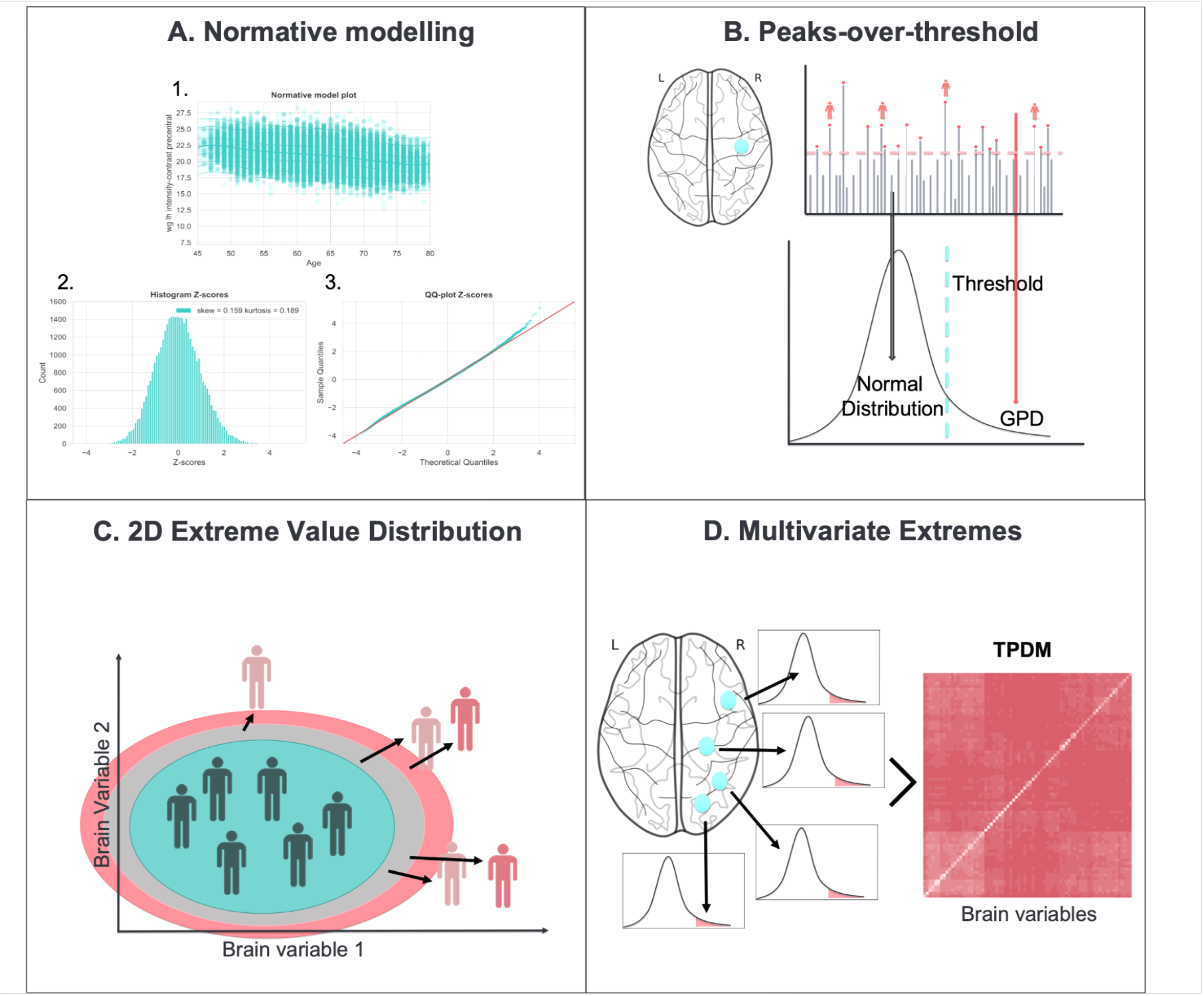
Showing the different types of extreme value statistics that can be used with normative modelling. A. Normative model created for one IDP (27012) modelled as a function of age using a Bayesian Linear Regression (BLR) model. 1. Shows the normative model fit with the centiles of variation. 2. Shows the histogram of the deviation or z-scores. 3. Shows the quantile-quantile (QQ) plot of the z-scores. B. Extreme value statistics using the peaks over threshold method on the z-scores. In general, the bulk of the distribution is modelled by a normal distribution and the tails converge to a Generalized Pareto Distribution (GDP). C. The extreme value bivariate distribution for two dimensions. D. Modelling multivariate extremes for more than two variables. Here, we summarize the tail dependence with a so-called tail pairwise dependence matrix (TPDM), similar to a PCA for the bulk of the distribution.

However, for didactic purposes, we start by describing the block maxima approach. This involves dividing the data into blocks of a certain size and then finding the maximum value in each block. For instance, in hydrology blocks are usually taken to be years, and the annual maximum is measured across each block. The goal is then to fit a statistical model to estimate the probability of these maxima.

More formally, we consider the deviation scores derived from the normative model. These are denoted *z*, which are realizations of the random variable *Z*. We use uppercase letters for random quantities and lowercase letters for actual measured values to differentiate between the two. We then consider a set of *n* independent variables *Z*_*n*_, having a common distribution function *F* and aim to model the statistical behaviour of the maxima:

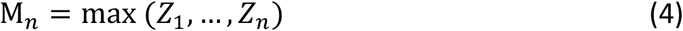

Usually, under the block-maxima approach *Z*_*n*_ are grouped according to a logical block unit or time frame, such as a time-block in fMRI, one full scanning session for MRI, or another repeated time measure in hours, months, and or years. Alternatively, one can also define a block in the spatial domain such as a region of interest (ROI) in the brain (or in our case one IDP). We then aim to determine an appropriate probability distribution function *F* and fit this to observed data. It has been shown (38,39) that the probability distribution of maxima *M*_*n*_ converges to a generalized extreme value distribution (GEVD), when rescaled with parameters *b*_*n*_ and *a*_*n*_ is defined by:

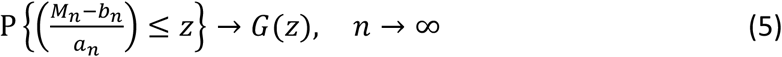

For constants *a*_*n*_ > 0, *b*_*n*_ ϵ *R* and *z* the upper end of the tails, we have

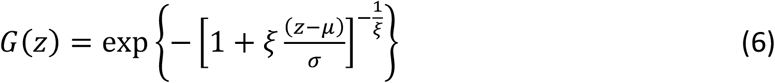

This probability distribution *G*(*z*) is defined by the shape *ξ*, location *µ* and scale *σ* parameters. The GEVD can be further separated into the Gumbel, Fréchet, and Weibull families, depending on the shape of the data generating distribution. Further details are provided in the supplementary material and in (7). What is remarkable about this result is that the GEVD is the only possible asymptotic limit for the distributions of *M*_*n*_, regardless of the form of the common data generating distribution *F*. Therefore, the theorem serves as an extreme value version of the central limit theorem (23).

The block maxima approach, although easy to employ, is not very efficient, because it ignores large parts of the data that could be relevant (for example, the second and third-highest values in each block which are potentially also informative about extreme behavior). Furthermore, this approach is mainly specified for data that have spatial or temporal block structure. Therefore, in this work, we employ the peaks over threshold approach. Instead of looking at the maximum value in each block, this method focuses on modelling values that exceed a certain threshold. A statistical model is then fit to these “peak” or extreme values. Mathematically, the peaks over threshold approach can be stated as:

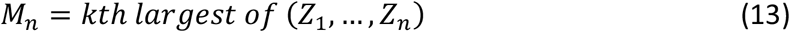

With *k* chosen to be an appropriately large threshold. In a similar manner to the convergence of the GEVD, the probability distribution of this model converges to a generalized Pareto distribution (GPD). This is derived by building on the results above, that is by assuming that *p*{M_*n*_ ≤ *z*} ≈ *G*(*z*). Then, by specifying that for a large enough threshold *k*, the distribution function of (*Z* − *k*), conditional on *Z* > *k*, is approximately:

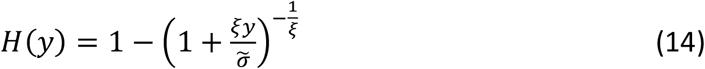

which is defined by the shape *ξ* and scale 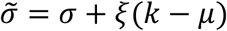 parameters. For more details see the supplement and (23).

### 2.3 Thresholding Procedures

Prior to computing the TPDM to capture the extremal dependence in the deviations from the normative model (i.e. the z-scores) it is necessary to transform the data and determine a suitable threshold, *k*, above which a sample is considered extreme. The threshold should not be too stringent as then there will be too few values to model the tails of the distribution accurately, but also not too low, as then the shape and scale parameters will be incorrect. One way to approach this problem is by using graphical tools with several thresholds and deciding on the best threshold based on their outcomes. This is the approach pursued here. In more detail, we follow the method proposed by (25) and transform the marginal distribution for each IDP, so that it fits the framework that requires z-scores to be regularly varying. Regular variation is a theoretical requirement underlying the statistics of extremes. In essence, if a random vector is multivariate regularly varying, it means that its joint tail decays according to a power function. This requirement is better satisfied after transforming the data. To achieve this, we transformed the marginal distribution of each variable using a rank transform. The rank order transformation smoothens effects of non-normality, outliers, unequal variances etc, but other transformations can be considered, see (34). Then, we applied the nonparametric marginal transformation proposed by 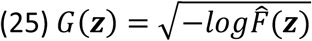, with 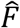 the original marginal distribution of the z-scores. This essentially ensures the marginal distributions of each variable has a distribution that is close to Fréchet. Next, to be able to threshold in multiple dimensions, we transformed the variables into polar coordinates (*r*, ***ω***) and selected the threshold using visual procedures outlined in the supplement.

One graphical tool that can be employed for the selection of an accurate threshold is the mean residual life plot (MRL-plot) (13), which involves plotting the value of a threshold, *k*, against the “mean excess”, i.e. the exceedances at *k*, minus *k*. Another graphical tool that can be employed is the parameter or threshold stability plot (40), which involves plotting model parameters as a function of the different thresholds. We look at different estimates of the parameters for the GDP and decide on an optimal threshold based on the lowest value where the plots are approximately stable. Examples of these plots are shown section 3.2 below.

Choosing the optimal (multivariate) threshold for the TPDM estimation includes all the same difficulties as selecting a threshold for univariate extreme value statistics. Commonly, it is selected with a visual inspection of all the graphs presented in the univariate result section. However, when the dimensions become large this becomes infeasible. Therefore, we follow the suggestions given in (25,26) we advocate selecting several variables manually. This is the approach we follow in this manuscript and several examples are shown in the results and supplement.

### 2.5 Framework for the tail pairwise dependence structure

With univariate extreme value statistics, we can model the tails in one variable. However, in many circumstances, we are interested in multiple variables simultaneously, which calls for a multivariate approach. For example, when there is a strong tail dependence between certain locations in the brain or between different imaging phenotypes derived from the data. This can be done using multivariate extremes, where our objective is to characterize and understand the dependence structure in the far upper joint tail, see Figure 3. There are several methods to model the multivariate joint tails, for a full overview see (22). In the bivariate case, we can fit a bivariate extreme value distribution, which tries to estimate the probability of the variable being large whilst the other is also large, Figure 3. C, however, this approach is very difficult to scale up to a higher dimension. In our case, the chosen multivariate extreme value method is based on PCA (25), called the tail pairwise dependence matrix (TPDM), see Figure 3. D.

As for the univariate method, the method used here for understanding the tail pairwise dependence structure is based on the framework of regular variation. We therefore assume ***Z*** is a regularly varying random vector. In general, for multivariate regular variation it is convenient to consider the variables in polar coordinates:

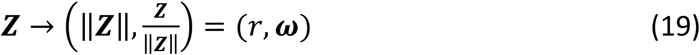

With **ω** ϵ ℝ^*N*^ a location on the unit sphere 𝕊 and *r* is the length of the vector (8). Then for a given *A*, a set of large values away from the origin, we can calculate the probability of ***Z*** as:

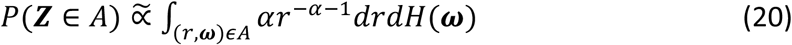

With 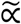 signifying “approximately proportional to”, *α* > 0, *r* the magnitude or radial component of the location, ***ω*** the location on the unit sphere 𝕊, and *H* the angular measure on the unit sphere 𝕊, see Figure 4. In the integrand, *r* follows a power-law distribution according to *α* the reciprocal of the shape parameter of the GPD(25,26).

**Figure 4.**
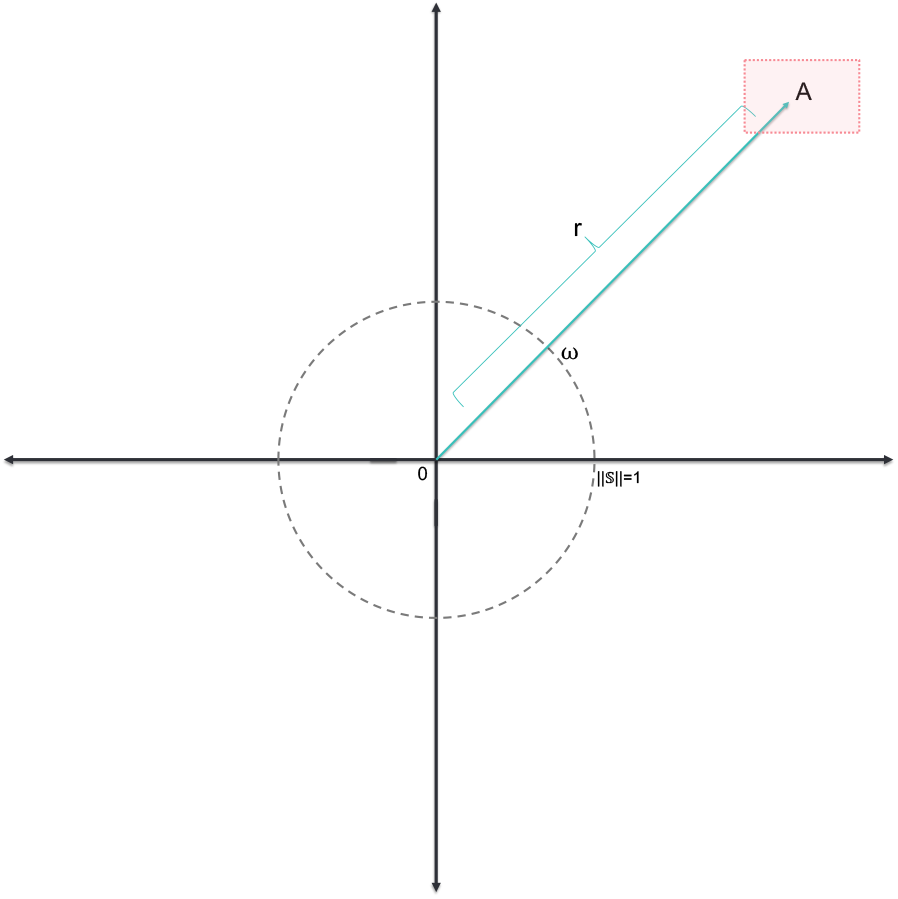
Illustration of the applied polar transformation of the data. the value indicated by the radial component r becomes independent from the angular component ω. Adapted from (9).

### 2.6 Tail pairwise dependence matrix

The angular measure *H* in lower dimensions can be modelled parametrically. However, as the dimension size increases, there is insufficient information in the extremes to fully capture *H*. Instead of fully modelling the dependence structure contained in *H* an approximation can be made with the tail pairwise dependency matrix (TPDM). If we assume ***Z*** to be a regularly varying random vector with a common tail index *α* and angular measure *H*. Then each element in the TPDM, Σ_***z***_, is specified according to:

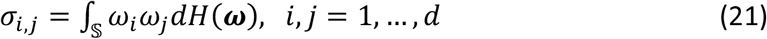

The TPDM has similar interpretation properties as a standard covariance matrix only specified for the extremes (25). Thus, if two variables are independent in the outer tails *σ*_*i,j*_ = 0 and the diagonal element *σ*_*i,j*_ = *b*^2^, with *b* the scale. The TPDM is positive definite and symmetric and therefore the eigenvalues are positive and the eigenvalues real (25,26).

### 2.7 Estimation of the TPDM

We estimated the dependence in the tails via the TPDM for ℤ = ℝ^*N*×*D*^, which is a matrix containing all data for *D* IDPs and *N* subjects. For this framework, the variables should be heavy-tailed with a common tail index *α* = 2. The framework for regular variation outlined earlier necessitates that all variables exhibit heavy-tailed behaviour, with a shared tail index denoted by *α*. To ensure this assumption holds, we first transformed the marginals, which is a common procedure in extreme value statistics (8). We first used a rank order transformation to obtain 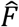, the estimated marginal cumulative distribution function (CDF) from the data. The rank order transformation smoothens effects of non-normality, outliers, unequal variances etc, but other transformations can be considered, see (34). Then we transformed our 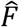 with 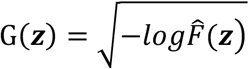 the CDF of a Fréchet random variable (25). Afterward, a polar coordinate transformation was applied and a threshold of the 95^th^ quantile was chosen for the kth largest radius, see Figure 4. Finally, we estimated the TPDM according to:

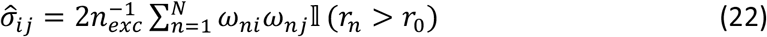

Where *r*_0_ the threshold for the radial component and *n*_*exc*_ the number of exceedances (i.e. observations that pass the threshold).

### 2.8 Extreme principal components analysis

Next, we perform an eigendecomposition (25) of the TPDM or 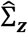 to obtain the eigenvectors (i.e. extreme PCs). This yields an orthonormal basis 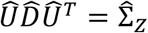, with *D* the diagonal matrix of eigenvalues (*λ*_1_, …, *λ*_*l*_) and *U* = (*u*_1_, …, *u*_*l*_) the matrix of eigenvectors.

Two of the goals of standard covariance PCA are to extract the most important information and to analyze the structure of this information in terms of the original variables (35). To achieve the first goal, we select the top eigenvectors by ordering the eigenvalues that explain the largest ‘scale’, which is the extremal equivalent of variance in conventional (covariance) PCA. This is therefore similar to ordering standard eigenvalues which explain the most variance for covariance PCA. To achieve the second goal, i.e. to understand the contribution of each variable to the extreme PCs, we compute loadings to understand the basis we derived in terms of its original variables by estimating the individual contributions. We define the loadings for each component 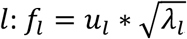, *w*ith *f* the loading, *u* the eigenvector and *λ* the eigenvalue. The contribution of each IDP *d* to the component *l* is then calculated by: 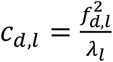, with *λ*_*l*_ the eigenvalue of the *l*-th component (35).

### 2.9 Evaluation

Finally, we correlate the extreme PCs with all the behavioural phenotypes according to the FUNPACK categorization described in section 2.1 above. We achieve this using pointwise biserial correlation or Spearman correlation depending on whether the phenotype is continuous or categorical for the leading principal components, followed by Bonferroni correction for multiple comparisons across nIDPs and PCs. We also perform a standard correlation PCA approach as a comparison. However, we emphasise that these two approaches are complementary as one is specified for the tails of the distribution and the other is specified for the tails of the distribution. Therefore, they yield different information, as we will show in the results.

## 3. Results

### 3.1 Univariate extreme value results

In this section, we demonstrate the peak-over-threshold method for one IDP and show how an optimal threshold can be decided and the tails accurately modelled. In Figure 5, we show the different possible thresholds for one IDP, volume of white matter hypointensities (WMH; UKB field ID 26528). This is a suitable choice because it is the most highly non-Gaussian phenotype we model and is therefore most challenging variable for our pipeline (however, please note that after fitting the normative model, the z-scores should be closer to Gaussian). This IDP is shown in Figure 5 below.

**Figure 5.**
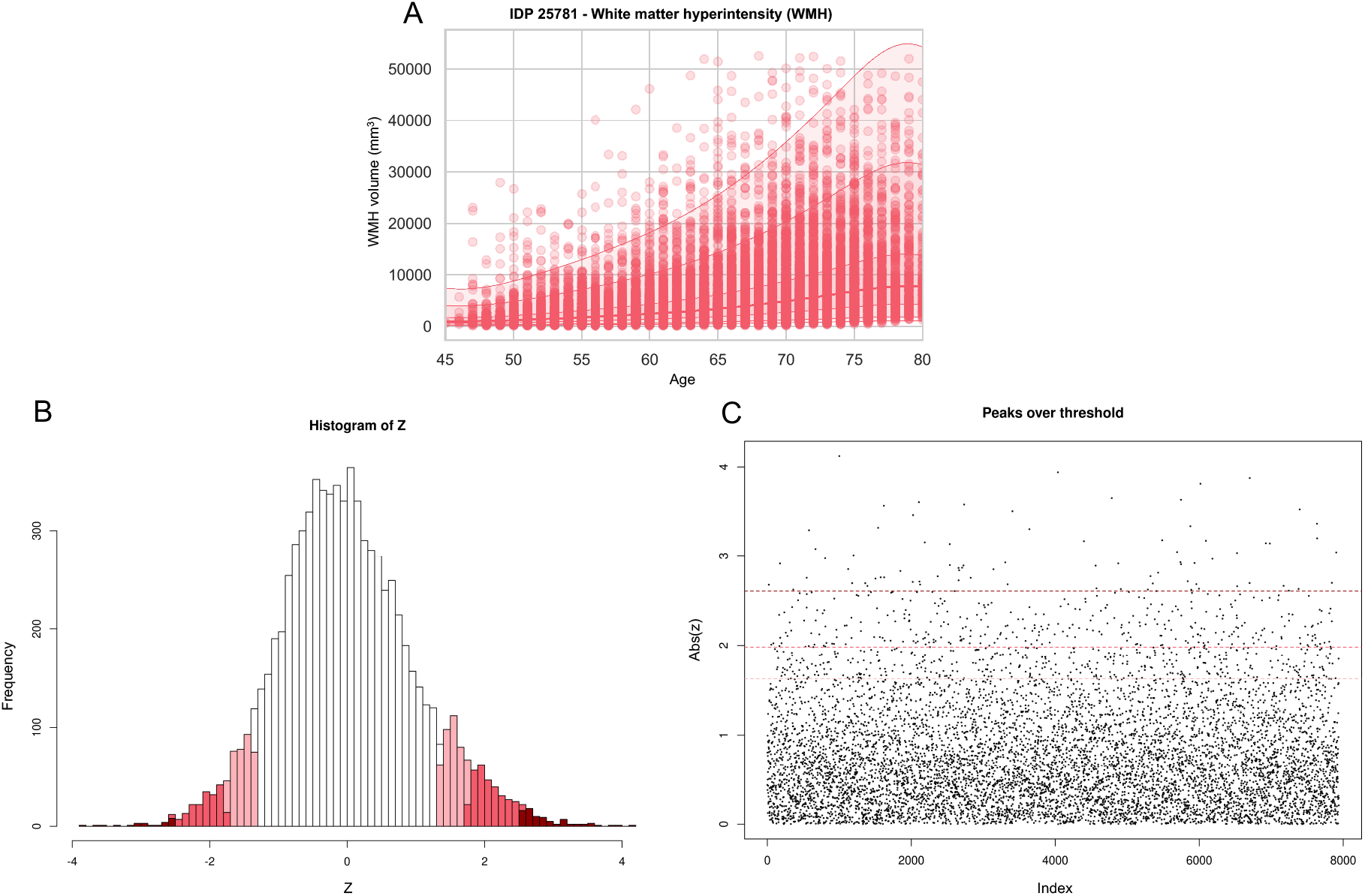
Showing the univariate peak over threshold method for one IDP. A. Warped BLR normative model result for the white matter hyperintensity IDP, demonstrating the mean and centiles of variation. B. Histogram of the z-scores showing different thresholds that could be employed in the peaks over threshold method (90th, 95th and 99^th^ percentiles). C. Scatterplot for the peaks over threshold method demonstrating three possible thresholds of the 90th, 95th and 99th percentiles. We see that the sparsity of the points increases with higher threshold values.

To decide between the different thresholds, we used graphical tools such as the mean residual life plot and shape and scale plots. In Figure 6, we present the mean residual life plot for the WMH phenotype. To help interpret this plot, it is assumed that the threshold should be placed where the plot starts diverging from the linear trend. In this case, we can observe that the plot is relatively linear until the 0.95 mark.

**Figure 6.**
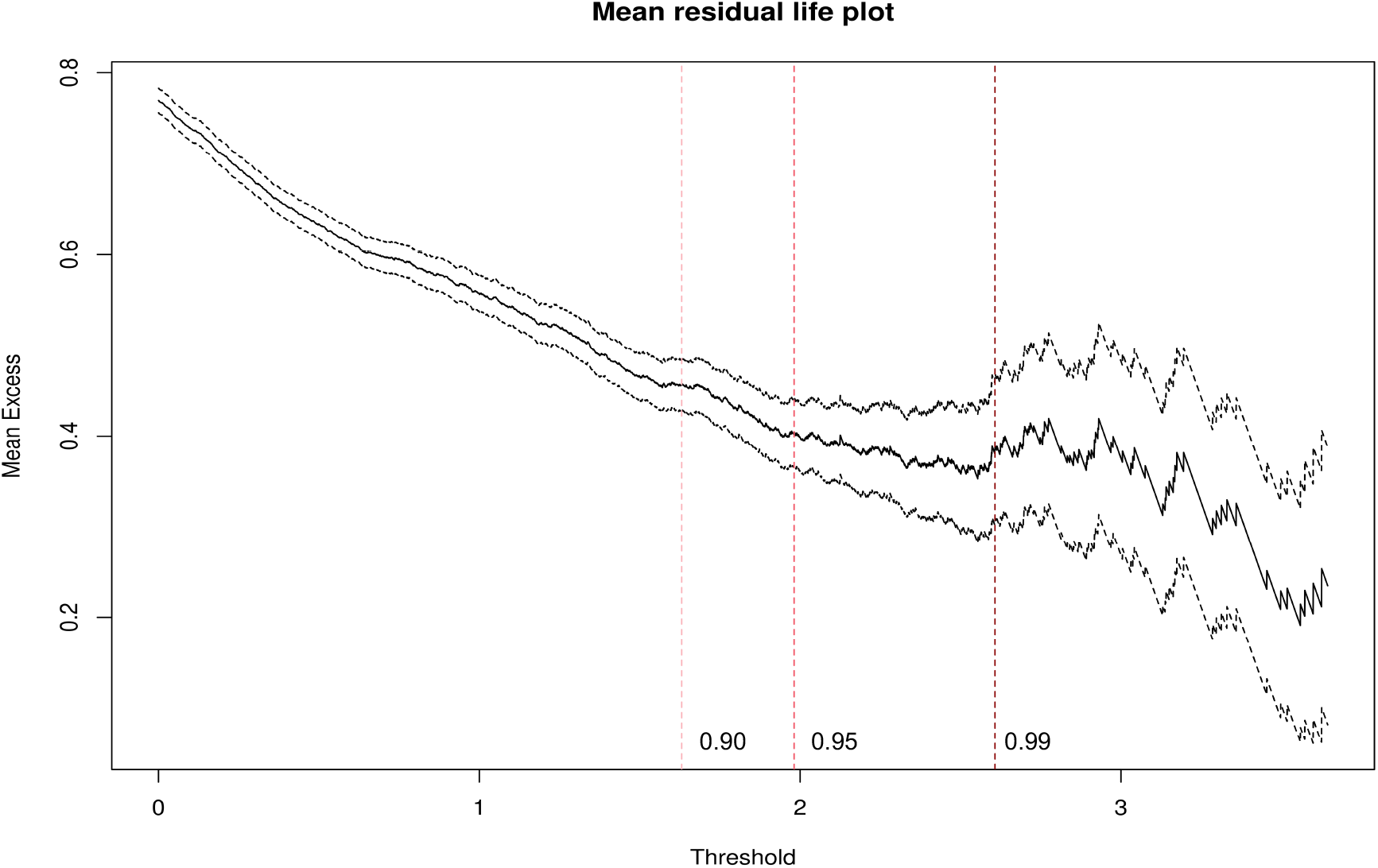
Showing the mean residual life plot of the WMH IDP data. We can observe that the plot is relatively linear up to the 0.95 threshold, indicating that this could be a good threshold.

In Figure 7, we show the parameter threshold stability plot of the modified scale and shape parameters. The dots demonstrate the scale and shape parameters for the GPD at different thresholds. The lines show the 95% confidence interval. Again, this makes it clear that the threshold should be around the 0.9 to 0.95 mark.

**Figure 7.**
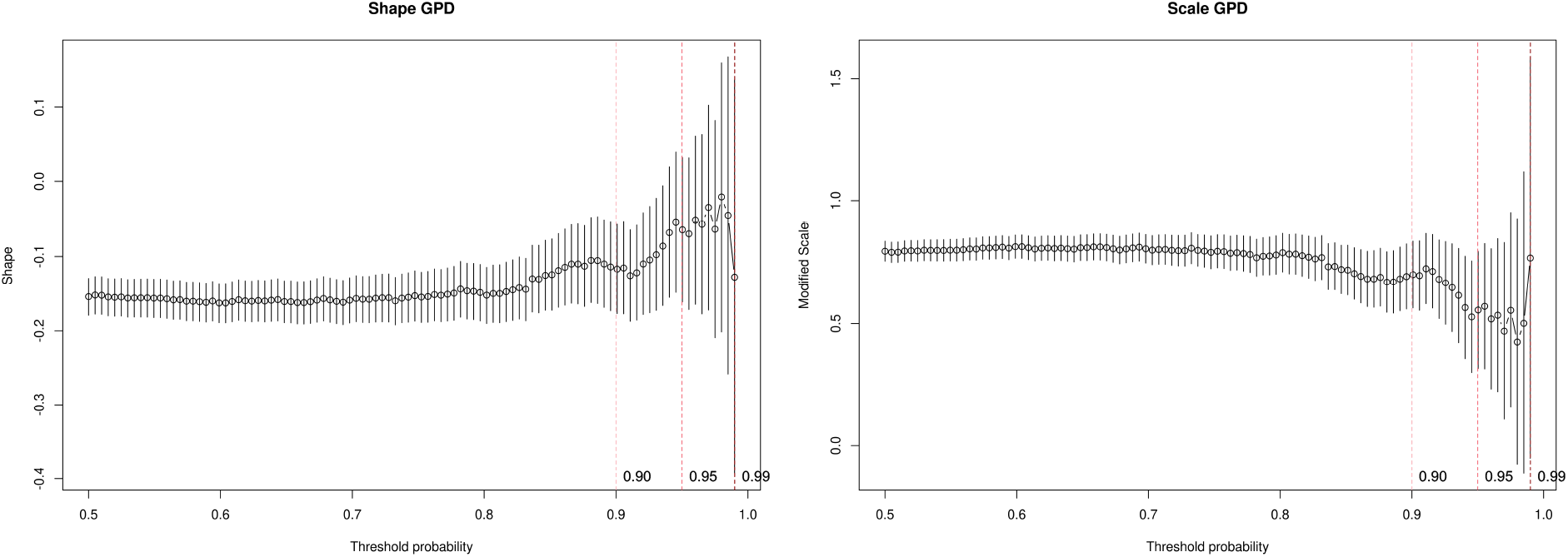
Showing the parameter threshold stability plot of the modified scale and shape parameters. The dots demonstrate the scale and shape parameters for the generalized Pareto distribution (GPD) at different thresholds. The lines show the 95% confidence interval. An optimal threshold is chosen at the point where the parameters are both relatively stable in this case around the 95th percentile.

On the basis of these techniques, we set the threshold we set the threshold as the 95^th^ quantile, i.e. *r*_0_ = 247. This yields *n*_*exc*_ = 1960 values exceeding this threshold, which is low enough to give us sufficient data and high enough to estimate the GPD parameters accurately. Once the threshold is decided, we can estimate the parameters using maximum likelihood optimization. We then optimized the profile log-likelihood to decide the scale and shape parameters for the generalized Pareto distribution. The more peaked the log-likelihood is the higher our confidence in our estimates. In the supplement, we show several model diagnostic plots, such as the QQ-plot and return level plot, yielding straight plots. The straight line indicates that the fitted function is appropriate for the data (34). The final scale and shape parameter estimates are: threshold: 1.9613, exceedances: 397, scale estimate 0.425 ± 0.03 (SE), shape estimate -0.065 ± 0.05 (SE).

### 3.2 Multivariate extreme value results

#### 3.2.1 Thresholding

As discussed in the methods, to be able to threshold in multiple dimensions, we transformed the variables into polar coordinates (*r*, ***ω***), then use multiple diagnostic plots to select the threshold, *r*_0_. Several examples are shown in supplementary Figures 7, 8 and 9. On the basis of these plots, we decided the 0.95 quantile was a good common threshold above which the examined parameters seemed relatively constant.

#### 3.2.1 Extreme principal components analysis

After computing the TPDM and its eigendecomposition, we show the scree plot of the scale in Figure 8 below. Briefly, the first extreme PC explains 17.64% of the scale and the second extreme PC explains 3.71% of the scale. We therefore selected the first two extreme PCs, as our informative basis.

**Figure 8.**
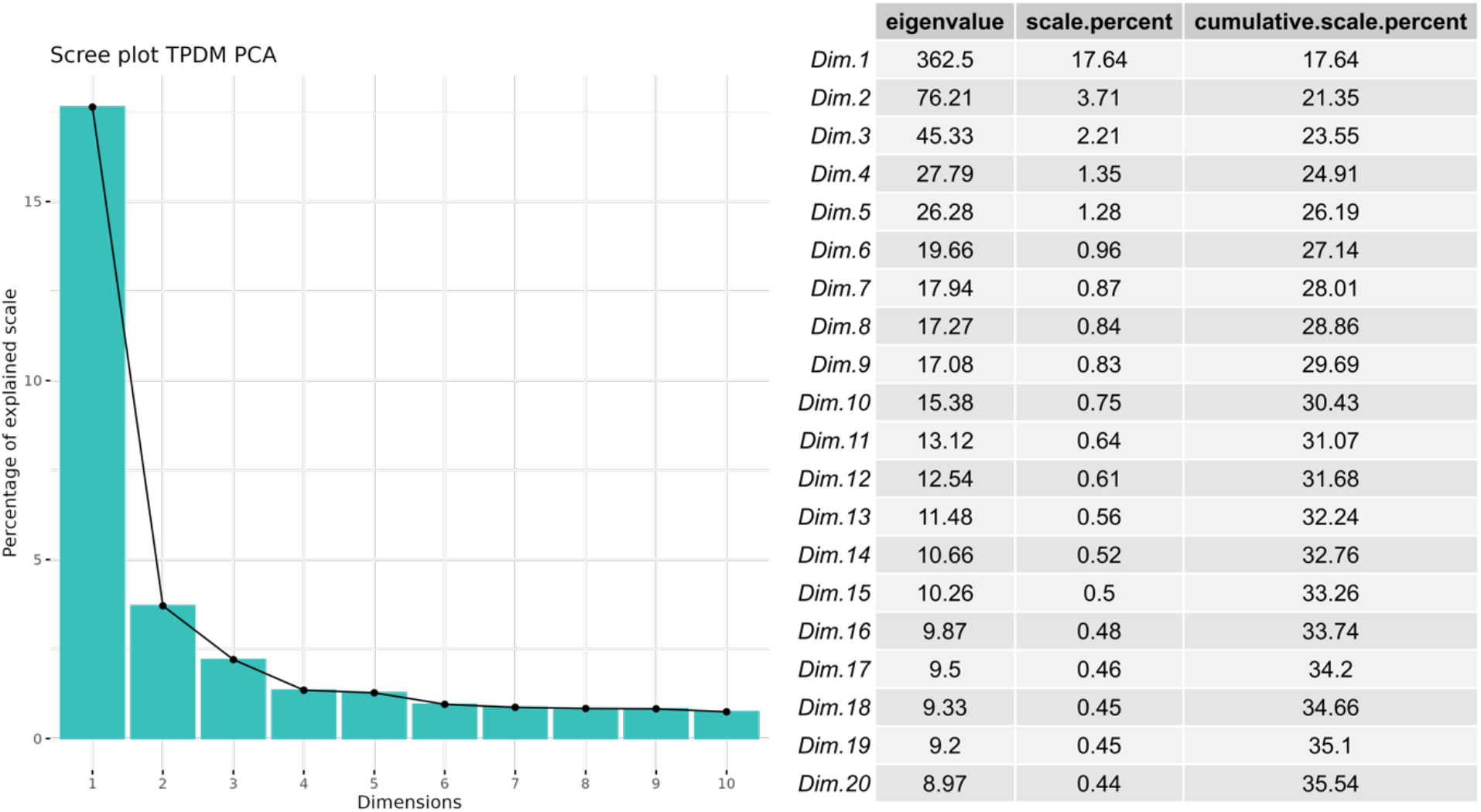
Showing the scree plot and table of scale values of the eigenvalue decomposition for the TPDM matrix. We can see that a large percentage of the scale is explained by the first extreme component.

In Figure 9, we plot the highest contributions (i.e. loadings) of the IDPs to the first two extreme PCs. A line is added to denote the average contribution (1/*l*). The first extreme PC is dominated by regional and tissue volume IDPs and cortical area IDPs and the second extreme PC is dominated by white matter tract diffusivity IDPs, showing that in the extreme tails these variables vary the most together and explain the largest amount of scale for all the IDP data. This shows that many phenotypes have very strong tail dependence, which underscores the need for a multivariate approach.

**Figure 9.**
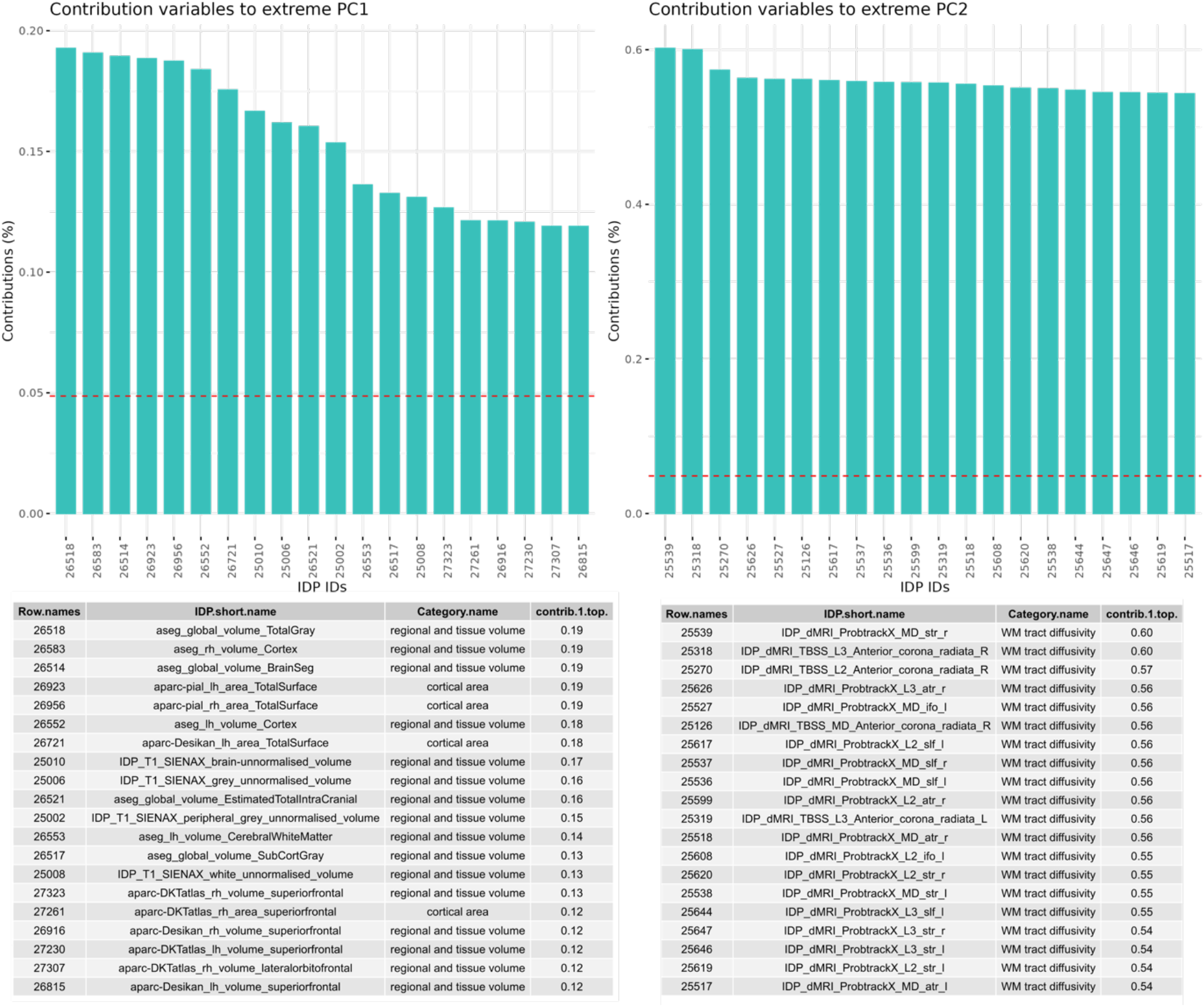
List of the top 20 contributions of the different IDPs to the first and second extreme PCs. We can see that the first extreme PC is largely dominated by IDPs relating to whole brain tissue volume and area. The second extreme PC is largely dominated by IDPs relating to white matter tract diffusivity.

For comparison, we show the comparable scree plots and loadings for standard PCA in Supplementary Figures 10 and 11. While it is difficult to compare these approaches since they measure different things, we select the top two components in each case for subsequent analysis, which explain approximately 20% of scale or variance respectively for extreme or standard PCA.

**Figure 10.**
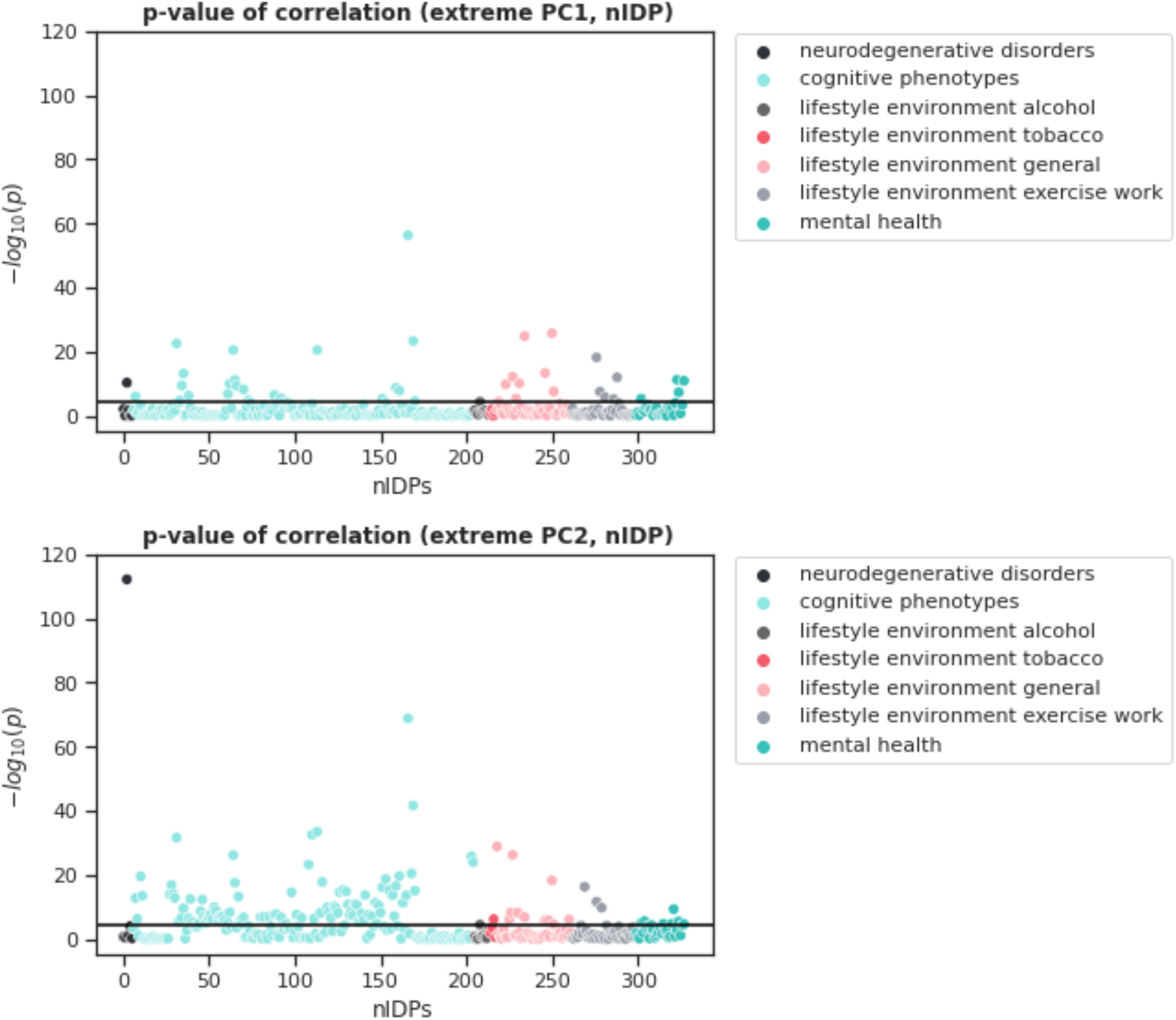
Showing two Manhattan plots of the log p-values for the Spearman correlation between the nIDPs and the first two extreme PCs. The black line demonstrates the Bonferroni-corrected p-value threshold. Thus, values passing this line indicate a significant PC-nIDP correlation. We can see that mostly cognitive measures and general lifestyle nIDPs correlate with the first two extreme PCs.

#### 3.2.2 Extreme PCs correlation with behavior

We next aim to correlate the extreme PCs to behavioral variables to ensure that the model can be used for behavioral predictions in clinical samples. In Figure 10, we present Manhattan plots of the p-values for the univariate correlation between the nIDPs and the first two extreme PCs. Across both PCs we detected 137 significant Bonferroni corrected associations. For standard PCA, we detected 158 Bonferroni corrected associations across the first two components. However, we emphasize that the number of detected associations is not the most important point of comparison, particularly recalling that extreme PCA is estimated on the basis of 20 times fewer data points (i.e. only samples exceeding the 95% threshold). Rather, we wish to emphasize that the associations detected are different between extreme and standard PCA, indicating that they capture different information. This can be seen by comparing Table 1 (extreme PCA) with supplementary Table 1 (standard PCA), both of which show the aggregate effects across the first two components. More specifically, extreme detect associations with phenotypes that could arguably be expected to have a more extreme effect on brain structure or function. For example, associations are detected between extreme PCs and multiple sclerosis and alcohol use, but not with standard PCs. In contrast, standard PCA detects stronger associations with general environmental factors. Note also, that this is not just a thresholding effect and the pattern of effects remains even at a more lenient threshold (not shown).

**Table 1.** Top 10 associations between non-imaging derived phenotypes (nIDPs) and extreme principal components, grouped per category. The p-values reported in the table are the maximum p-value across the first two components. Only associations surviving Bonferroni correction across nIDPs and components are reported.

| UKB IDP ID | $-\log_{10}(p)$ | Description |
| --- | --- | --- |
| **Neurodegenerative disorders** |  |  |
| 20002-0.1261 | 112.2 | Multiple sclerosis |
| **Cognition** |  |  |
| 20016-2.0 | 68.8 | Fluid intelligence score |
| 20128-2.0 | 41.7 | Number of fluid intelligence questions attempted within time limit |
| 6373-2.0 | 33.5 | Number of puzzles correctly solved |
| 6350-2.0 | 32.5 | Duration to complete alphanumeric path (trail #2) |
| 630-2.0 | 31.6 | Touchscreen duration |
| 4282-2.0 | 26.2 | Maximum digits remembered correctly |
| 23323-2.0 | 25.7 | Number of symbol digit matches attempted |
| 23324-2.0 | 24.0 | Number of symbol digit matches made correctly |
| 6348-2.0 | 23.3 | Duration to complete numeric path (trail #1) |
| 20023-2.0 | 20.5 | Mean time to correctly identify matches |
| **Lifestyle/environment alcohol** |  |  |
| 1588-2.0 | 4.5 | Average weekly beer plus cider intake |
| **Lifestyle/environment tobacco** |  |  |
| 20116-2.0 | 6.2 | Smoking status |
| **Lifestyle/environment general** |  |  |
| 3-2.0 | 28.9 | Verbal interview duration |
| 738-2.0 | 26.2 | Average total household income before tax |
| 6138-2.0 | 25.7 | Qualifications |
| 2139-2.0 | 24.8 | Age first had sexual intercourse |
| 3526-2.0 | 13.4 | Mother's age at death |
| 1835-2.0 | 10.1 | Mother still alive |
| 680-2.0 | 9.8 | Own or rent accommodation lived in |
| 728-2.0 | 8.3 | Number of vehicles in household |
| 1807-2.0 | 8.2 | Father's age at death |
| 6142-2.0 | 7.6 | Current employment status |
| **Lifestyle/environment exercise/work** |  |  |
| 1070-2.0 | 18.2 | Time spent watching television (TV) |
| 924-2.0 | 16.3 | Usual walking pace |
| 1190-2.0 | 12.0 | Nap during day |
| 1100-2.0 | 9.7 | Drive faster than motorway speed limit |
| 1090-2.0 | 7.5 | Time spent driving |
| 1120-2.0 | 5.7 | Weekly usage of mobile phone in last 3 months |
| 1170-2.0 | 5.2 | Getting up in morning |
| 1130-2.0 | 4.2 | Hands-free device/speakerphone use with mobile phone in last 3 month |
| 904-2.0 | 4.2 | Number of days/week of vigorous physical activity 10+ minutes |
| **Mental health** |  |  |
| 4570-2.0 | 11.3 | Friendships satisfaction |
| 4653-2.0 | 10.9 | Ever highly irritable/argumentative for 2 days |
| 4548-2.0 | 9.3 | Health satisfaction |
| 4581-2.0 | 7.3 | Financial situation satisfaction |
| 1970-2.0 | 5.6 | Nervous feelings |
| 1950-2.0 | 5.3 | Sensitivity / hurt feelings |
| 4537-2.0 | 4.8 | Work/job satisfaction |
| 2080-2.0 | 4.7 | Frequency of tiredness / lethargy in last 2 weeks |
| 2010-2.0 | 4.4 | Suffer from 'nerves' |
| 4631-2.0 | 4.2 | Ever unenthusiastic/disinterested for a whole week |

## Discussion

In this paper, we present a framework to characterize multivariate extreme tail behavior of brain imaging data. This is predicated on re-conceptualizing psychopathology as deviations from a normative pattern of functioning, where such deviations may be spread across multiple measures and have a complex correlation structure, although we consider that this framework will also be useful for many other neuroimaging applications where understanding tail behaviour is of interest. For example, our approach is naturally suited to anomaly detection applications where quantifying the statistical ‘surprise’ of a potential outlier is important, and indeed our approach is useful in any application where it is important to accurately quantify risk in the tails of a given distribution. We introduced the key notions underlying extreme value statistics, and outlined approaches suitable for fitting these models to neuroimaging data including heuristics for univariate and multivariate threshold selection. In order to accommodate tail dependency across multiple variables, we propose to apply an eigendecomposition to the tail pairwise dependency matrix (TPDM), yielding an extremal variant of PCA that can provide a good basis to further understand multivariate tail dependence. We show that this approach is useful to identify participants deviating from the normative model in multivariate space and that this meaningfully relates to clinical and demographic variables. Finally, we show that this extreme PCA approach detects fundamentally different (and complementary) information to the standard covariance PCA approach used in the field.

We applied this approach to normative models, which are now widely used to capture and understand the heterogeneity in the population by modelling individual deviations from a reference cohort. Although many applications have considered the extreme deviations from a normative model as their primary endpoints (4–6,16,17), a principled treatment of how to study extreme deviations from normative models has been lacking, which we now provide. In this study, we show that fitting an extreme value distribution to these deviations allows us to statistically quantify the magnitude of extremes (e.g. as quantiles of an assumed extreme value distribution) and to explore extreme tail behavior across multiple variables by summarizing extremal dependence structure in a TPDM. Previous research in other domains (e.g. meteorology, hydrology, climate and finance) has attempted to measure tail dependence between variables, as seen in (22). Our results show that the TPDM method is an effective tool for this purpose in neuroimaging data. The TPDM is similar to a covariance matrix (25), a well-established tool in neuroimaging research for understanding the relationships between different brain areas and modalities. However, the covariance matrix only captures variance around the mean, which does not fully correspond to tail dependence.

We demonstrated how to implement the multivariate extreme value analysis on a large dataset from the UK Biobank, using image-derived phenotypes (IDPs). In this application, we showed that the scale of the TPDM can be summarized by a low number of extreme principal components and that these can be meaningfully related to behavioural phenotypes, providing complementary information to a standard PCA. The multivariate extreme method presented in this paper could offer an alternative to methods mostly focusing on mean deviation scores. Although we focus on normative models here, the extreme value statistics approach is widely useful in neuroimaging, although we point out that fitting imaging data to normative models first is a natural preliminary step and provides important benefits. For example, it is easier to find a common threshold for multivariate analysis if data are first transformed via normative models to a common reference scale (i.e. Z scores). In contrast, it may be more difficult to do this if the raw data values are used which may have different units and scaling (e.g. combining cortical thickness estimates with subcortical volumes).

As with any statistical method, for extreme value statistics, one major consideration is that users should carefully consider the questions they want the model to answer. The extreme value statistics approach assumes *a priori* that there is more information of interest in the tails than in the bulk of the distribution. If users have a question pertaining to the general population and ‘healthy’ (or ‘average’) functioning of the brain, an extreme value approach would probably not be the most optimal. For these questions, classical approaches that focus on central tendency could be more appropriate. Therefore, we recommend users carefully consider the nature of their question and asses if it is focused on the mean or the tails of the distribution. The extreme value method is designed to give a more sensitive analysis of the multivariate tails of the distribution and is especially beneficial in applications where accurate quantification of risk is essential. This is, to a certain degree, shown in the analysis we conducted within UK Biobank, where for example more severe clinical phenotypes (such as multiple sclerosis) were more strongly associated with the extreme PCs than standard PCs.

In this paper, we have considered the extreme value framework from the standpoint of considering mental disorders as falling at the extremum of the normal multivariate range, which we consider to be both natural and principled. This offers a powerful tool for uncovering new insights into the underlying relationships between different neuroimaging modalities and for discovering new biological markers of disease. We have also aimed to provide guidance specifically oriented toward neuroimaging for the many considerations that are important in implementing extreme value statistics in practice; for example, the choice of approach (block maximum, peaks over threshold), the choice of threshold parameter, how to transform variables to meet the assumptions underlying extreme value theory and how to model dependence between variables. However, there are many avenues for future work: first, it would be interesting to model tail dependence in a more structured way, for example using graphical models (22) or using stronger constraints such as modelling statistical independence in tail behaviour, as is done classically using independent components analysis. Another future avenue could be to design appropriate mixture models to combine a classical model to fit for the bulk of the distribution and an extreme value fit to the tails. This would allow accurate modelling of both the bulk and tail. However, in this case giving adequate weight to the extremes and the bulk of the distribution is important. Finally, other applications could consider assessing extremal dependence in the spatial domain rather than across subjects (e.g. comparing extreme deviations of different clusters within brain images) or combining extremes across both space and subjects.

In conclusion, we presented a comprehensive framework for understanding and visualizing the dependency structure in the tails of different neuroimaging variable distributions. The method presented here is general and can be applied to several types of neuroimaging data, including combining across modalities. Therefore, we expect that this method would be an excellent choice for many applications where extreme brain deviations and their underlying dependence could have a significant effect on extreme behavior, ultimately taking us further in the quest for individualized and precision medicine.

## Ethics and Data availability

Ethical approval for the acquisition of the UK Biobank data was provided by the UK Biobank consortium, and the data were accessed under application number 23668 (Marquand)

## Data and Code availability

Data are available under request from UK Biobank. Analysis code for the present manuscript are available at https://github.com/CharFraza/extremes_normative_modelling and the PCNtoolkit software toolbox used for normative modelling is available at https://github.com/amarquand/PCNtoolkit.

## Author Contributions

AM, CB: conceptualisation, funding, supervision. CF, MZ: data analysis. All authors: drafting and revising the manuscript

