## Supplemental figures 1-12 for "Using extreme value statistics to reconceptualize psychopathology as extreme deviations from a normative reference model"

### Supplementary Methods

#### Sample

For the distribution of the age range and the number of participants in our dataset, see Figure 1.

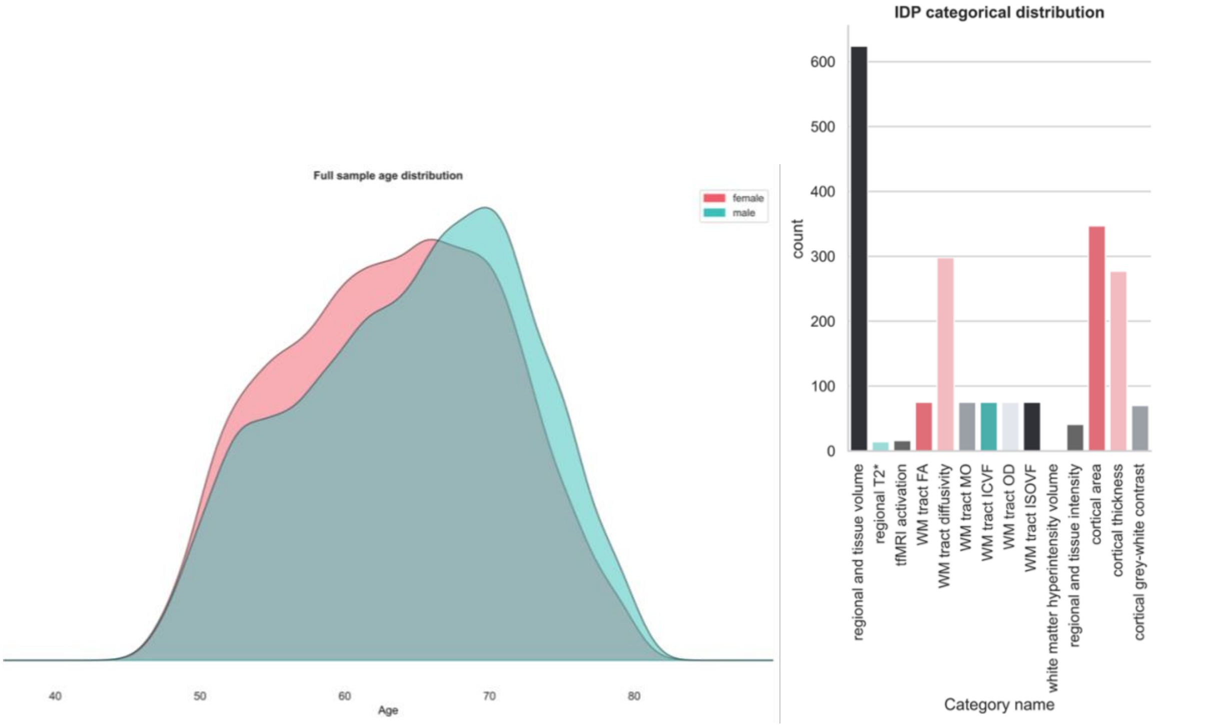


*Figure 1 – On the right - showing the age and sex distribution of the IDPs used in this study from the UK biobank. On the left - showing the distribution of the different IDP categories present in the UK Biobank dataset*

#### Normative model formulation

We estimated all the normative models using python 3.8.3 and the PCNtoolkit version 0.20. We performed a normative model with the Bayesian linear regression model (BLR) and likelihood warping. For a full explanation and details of the mathematical framework behind this model see (1). Here we will give a short overview of the mathematics behind the method. We define $\boldsymbol{y} = (y_{nd}) \in \mathbb{R}^{N \times D}$ with $y_{nd}$ the $d$-th IDP of the $n$-th subject. The covariates are collected into one matrix $\boldsymbol{x} = (x_{nm}) \in \mathbb{R}^{N \times M}$, where $x_{nm}$ is the $m$-th covariate of the $n$-th subject:

$$\boldsymbol{y} = \begin{bmatrix} y_{11} & y_{12} & \cdots & y_{1d} \\ y_{21} & y_{22} & \cdots & y_{2d} \\ \vdots & \vdots & \ddots & \vdots \\ y_{n1} & y_{n2} & \cdots & y_{nd} \end{bmatrix} \text{ and } \boldsymbol{x} = \begin{bmatrix} x_{11} & x_{12} & \cdots & x_{1m} \\ x_{21} & x_{22} & \cdots & x_{2m} \\ \vdots & \vdots & \ddots & \vdots \\ x_{n1} & x_{n2} & \cdots & x_{nm} \end{bmatrix}$$

We used the covariates age, sex and site. Here the sites are denoted by $s \in \{1, \dots, S\}$ with all subjects having the same sites. For each IDP we fitted a separate model. To keep the notation concise, we will concentrate on one specific IDP labelled $d$ and drop the subscript. Thus, for every IDP we denote $\boldsymbol{y} = (y_1, \dots, y_N)^T$ and take the set of independent variables $\boldsymbol{x}_n = (x_{n1}, \dots, x_{nM})^T$. For every subject we specified the model as follows:

$$\varphi(y_n) = \boldsymbol{w}^T \phi(\boldsymbol{x}_n) + \epsilon_s \quad (1)$$

Where, $\boldsymbol{w}^T$ is the estimated vector of weights and $\phi(\boldsymbol{x})$ is a basis expansion of the covariate vector $\boldsymbol{x_n}$. In our case, a cubic B-spline basis expansion with 5 evenly spaced knots was chosen. Empirically, this was enough to capture the curvature in space caused by the age covariate. $\epsilon = \mathcal{N}(0, \beta^{-1})$ is a Gaussian noise distribution with mean zero and noise precision term $\beta$ (the reciprocal of the variance). $\varphi(y_n)$ is a likelihood warping function used to accommodate non-Gaussianity of the residuals of the data in the original space. For the likelihood warping a SinhArcsinh function was employed:

$$\varphi(y_n, \boldsymbol{\gamma})_{SinhArcsinh} = \sinh\left(c * \operatorname{arcsinh}(y_n) - d\right) \quad (2)$$

With $\boldsymbol{\gamma} = (c, d)$ the identified parameters for the warping function. This method has been shown to be able to model Gaussian as well as non-Gaussian distributions (1). If non-Gaussianity is present in the data, there are other techniques besides likelihood warping that one can consider. A pre-transformation of the dependent variable, like a Box-Cox or log-transform, is one illustration. Finding the right transformation beforehand that is optimal across various datasets, however, can be quite difficult. The warped BLR model thus has the additional benefit of eliminating the extra step of selecting an appropriate transformation for each dataset by incorporating this in the model through likelihood warping (1,2). We captured the site variation using a fixed-effects model, according to (3). We performed the optimization using Powell's conjugate direction method by minimizing the negative log-likelihood. Afterward, the *z*-scores were calculated in the warped space for each subject, *n*, and IDP, *d*, in the test set as:

$$z_{nd} = \frac{y_{nd} - \hat{y}_{nd}}{\sqrt{\sigma_d^2 + (\sigma_*^2)_d}} \quad (3)$$

Where, $y_{nd}$ is the true response, $\hat{y}_{nd}$ is the predicted mean, $\sigma_d^2$ is the estimated noise variance (reflecting uncertainty in the data), and $(\sigma_*^2)_d$ is the variance attributed to modelling uncertainty, for the full derivations see (1,4). We evaluated the model fits according to several model criteria: explained variance ($R^2$), mean squared log-loss (MSLL), and skew and kurtosis. Together these criteria allowed us to assess the central tendency, performance of the warping function, as well as overall model fit. Afterward, we used the *z*-scores to estimate the extreme value distributions.

**Additional details about extreme value statistics**

Further details about the generalised extreme value (GEV) distribution and generalised Pareto distribution (GPD) are provided in the boxes below.

**The Generalized Extreme Value Distribution**

**Theorem 1:** If there exist sequences of constants $\{a_n > 0\}$ and $\{b\_n\}$ such that:

$$P\left\{\left(\frac{M_n - b_n}{a_n}\right) \le z\right\} \to G(z), \quad n \to \infty \tag{7}$$

Where $G$ is a non-degenerate distribution fucntion, then $G$ belongs to one of the following families:

$$\text{I: G(z)} = \exp\left\{-\exp\left[-\left(\frac{z-b}{a}\right)\right]\right\}, -\infty < z < \infty; \tag{8}$$

$$\text{II: G(z)} = \begin{cases} 0, & z \le b; \\ \exp\left\{-\left(\frac{z-b}{a}\right)^{-\alpha}\right\}, & z > b; \end{cases} \tag{9}$$

$$\text{III: G(z)} = \begin{cases} \exp\left\{-\left[-\left(\frac{z-b}{a}\right)^{\alpha}\right]\right\}, & z < b; \\ 1, & z \ge b; \end{cases} \tag{10}$$

for parameters $a$ > 0, b and, in the case of families II and III, $a > 0$. The extreme value distributions consist of three categories, which are commonly referred to as the Gumbel, Fréchet, and Weibull families, known as types I, II, and III, respectively. Each family is characterized by a location parameter (b) and a scale parameter (a). The Frechet and Weibull families also have a shape parameter (α). The Gumbel, Fréchet, and Weibull families can be merged into a unified family of models called the generalized extreme value (GEV) family of distributions.

**Theorem 1.1**: If there exist sequences of constants $\{a_n > 0\}$ and $\{b_n\}$ such that:

$$P\left\{\frac{M_n - b_n}{a_n} \le z\right\} \to G(z), \quad n \to \infty \tag{11}$$

For a non-degenerate distribution function $G$, then $G$ is a member of the GEV family:

$$G(z) = \exp\left\{-\left[1 + \xi\left(\frac{z-\mu}{\sigma}\right)\right]^{-\frac{1}{\xi}}\right\} \tag{12}$$

Defined on $\{z: 1 + \frac{\xi(z-\mu)}{\sigma} > 0\}$, where $-\infty < \mu < \infty,\ \sigma > 0$ and $-\infty < \xi < \infty$.

**The Generalized Pareto Distribution**

**Theorem 2:** Let $Z_1, Z_2, \dots, Z_n$ be a sequence of independent random variables with common distribution function $F$, and:

$$M_n = \max(Z_1, \dots, Z_n) \tag{15}$$

suppose that F satisfies Theorem 1.1, so that for large $n$,

$$\mathrm{P}\,\{M_n \le z\} \approx G(z) \tag{16}$$

Where

$$G(z) = \exp\left\{-\left[1 + \xi\left(\frac{z-\mu}{\sigma}\right)\right]^{-\frac{1}{\xi}}\right\} \tag{17}$$

For some $\mu, \sigma > 0$ and $\xi$. Then, for a large enough threshold $k$, the distribution function of $(Z - k)$, conditional on $Z > k$, is approximately:

$$H(y) = 1 - \left(1 + \frac{\xi y}{\tilde{\sigma}}\right)^{-\frac{1}{\xi}} \tag{18}$$

Defined on $\{y\colon y > 0\ and\ \left(1 + \frac{\xi y}{\tilde{\sigma}}\right) > 0\}$, where

$$\tilde{\sigma} = \sigma + \xi(k - \mu)$$

This family of distributions is called the generalized Pareto family.

### Supplementary Results

#### Bayesian Linear Regression model fit

We evaluated the performance of the BLR model and the likelihood warping by looking at the explained variance, MSLL, kurtosis, and skewness, see Figure 2.

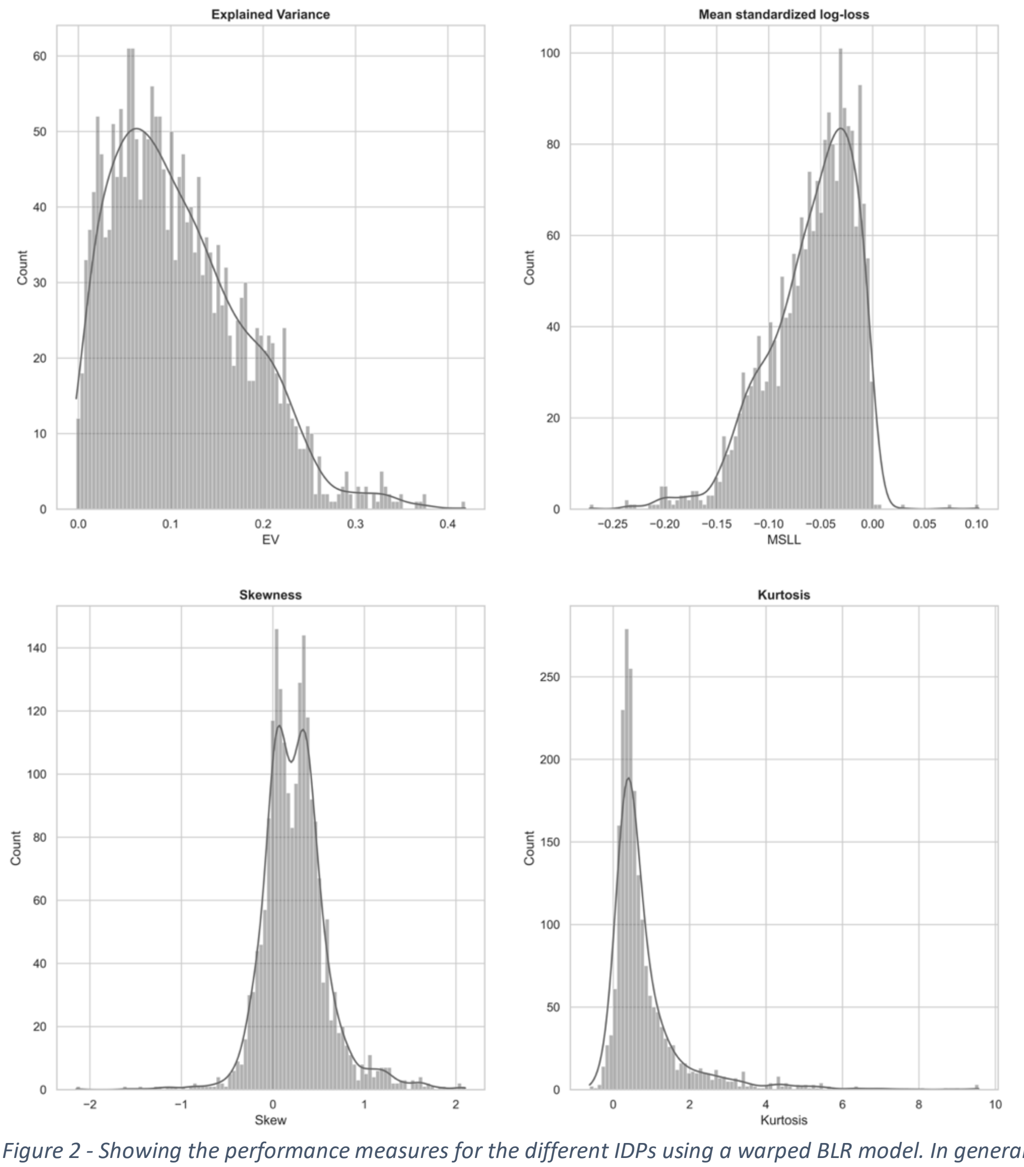


*Figure 2 - Showing the performance measures for the different IDPs using a warped BLR model. In general, for an optimal model fit the skew and kurtosis would both be distributed around zero.*

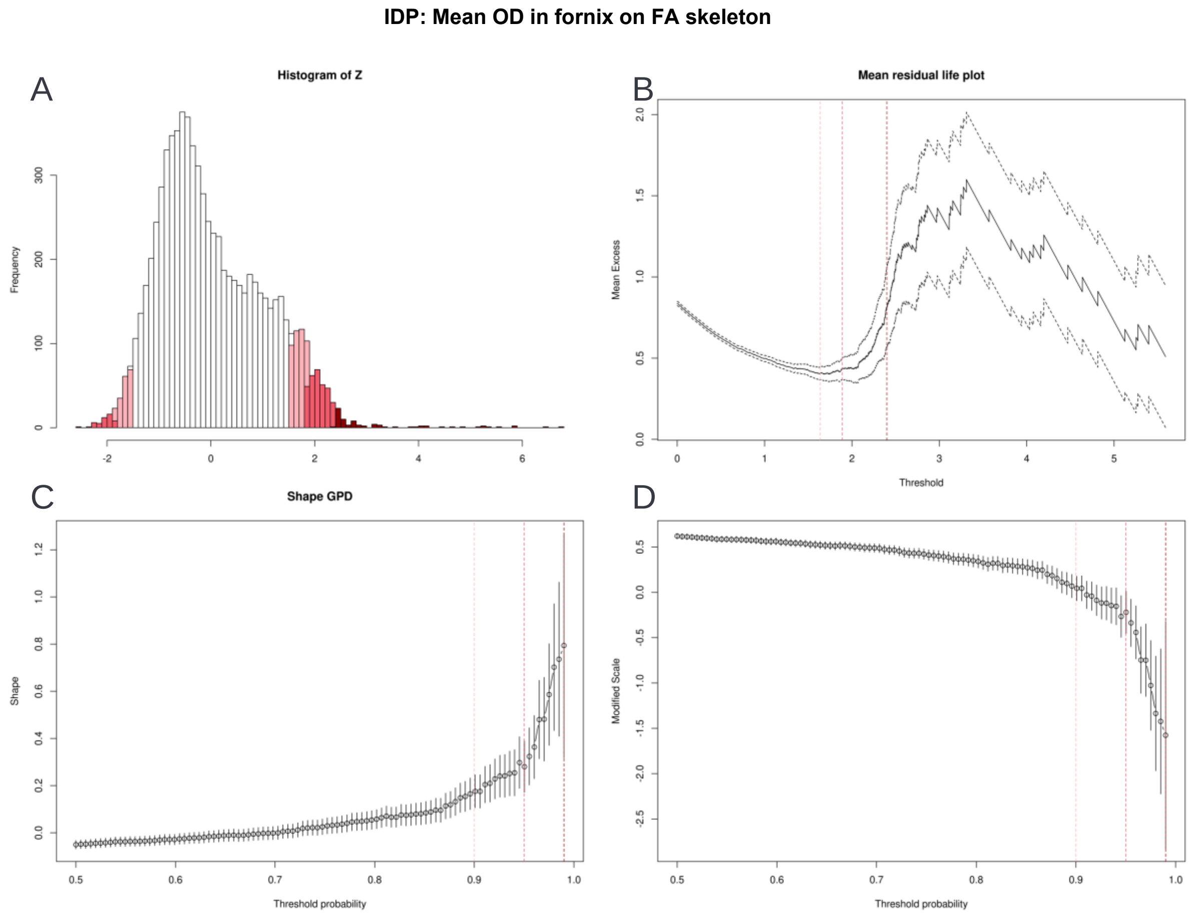


*Figure 3 - Displaying various threshold selection results for IDP 25397 - mean OD in fornix on FA skeleton. A. illustrates the histogram of the z-scores, demonstrating different thresholds that could be employed in the peaks over threshold method. B. displays the mean residual life plot, showing a relatively linear plot up to the 0.95 threshold, indicating that this could be a good threshold. C. and D. show the parameter threshold stability plot of the modified scale and shape parameters. The dots demonstrate the scale and shape parameters for the Generalized Pareto distribution (GPD) at different thresholds. The lines show the 95% confidence interval. An optimal threshold is chosen at the point where both parameters are relatively stable, in this case, around the 95th percentile.*

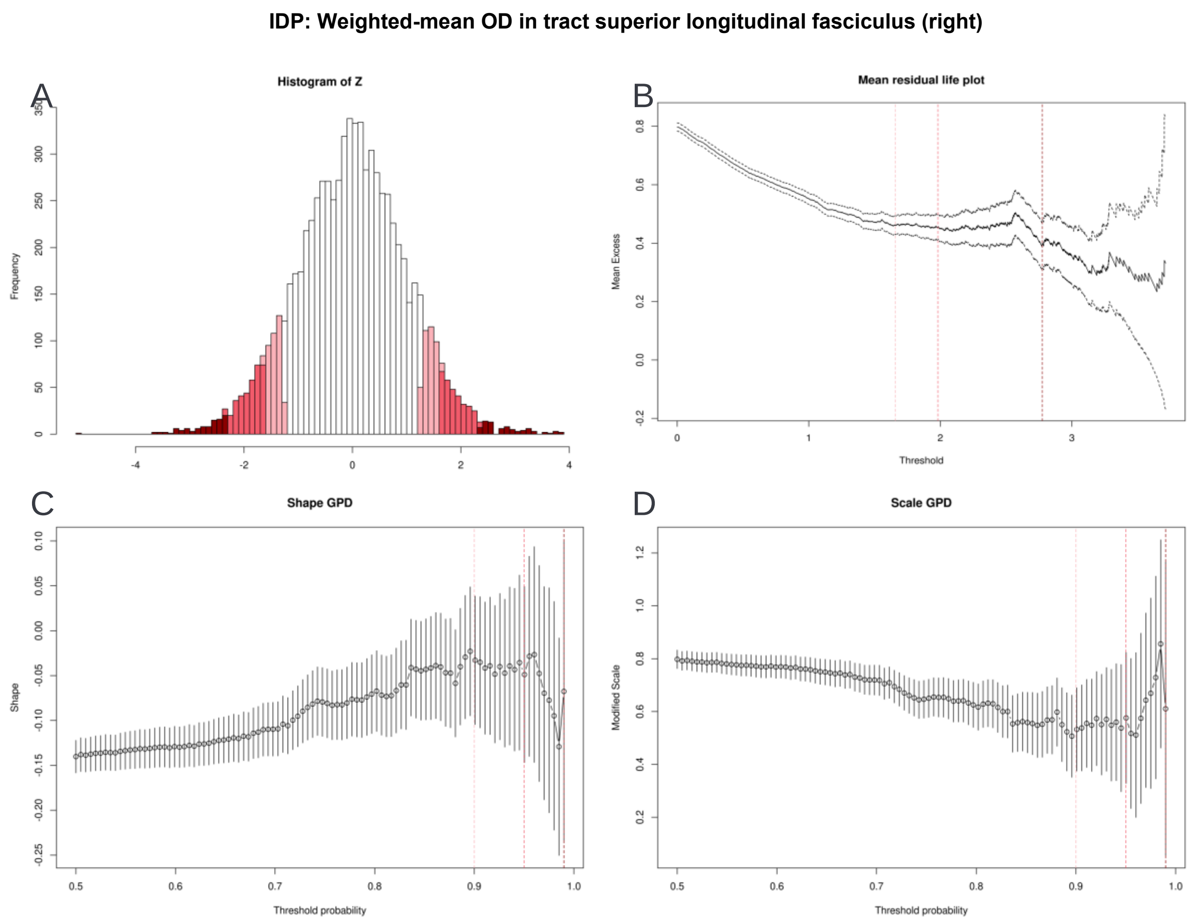


*Figure 4 - Displaying various threshold selection results for IDP 25699 - weighted-mean OD in tract superior longitudinal fasciculus (right). A. illustrates the histogram of the z-scores, demonstrating different thresholds that could be employed in the peaks over threshold method. B. displays the mean residual life plot, showing a relatively linear plot up to the 0.95 threshold, indicating that this could be a good threshold. C. and D. show the parameter threshold stability plot of the modified scale and shape parameters. The dots demonstrate the scale and shape parameters for the Generalized Pareto distribution (GPD) at different thresholds. The lines show the 95% confidence interval. An optimal threshold is chosen at the point where both parameters are relatively stable, in this case, around the 95th percentile.*

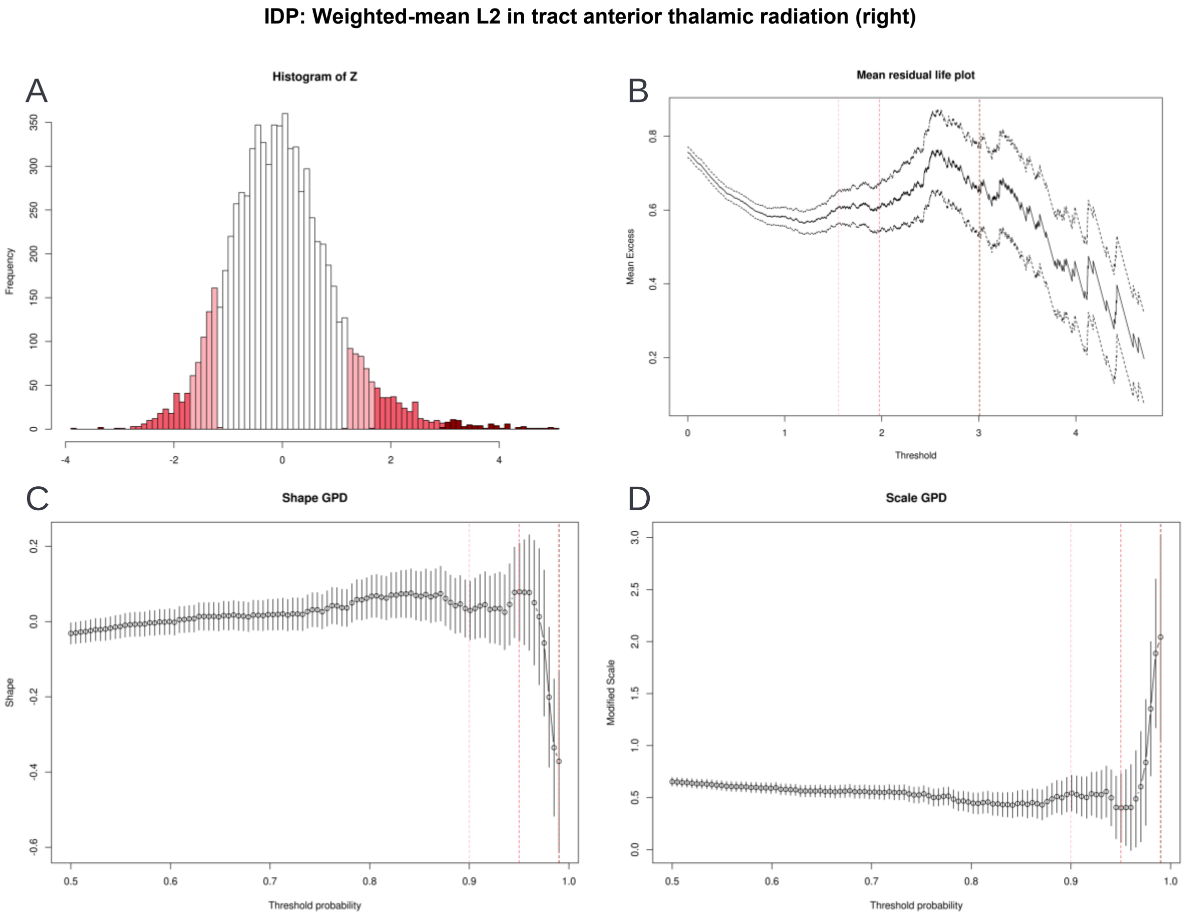


*Figure 5 - Displaying various threshold selection results for IDP 25599 - Weighted-mean L2 in tract anterior thalamic radiation (right). A. illustrates the histogram of the z-scores, demonstrating different thresholds that could be employed in the peaks over threshold method. B. displays the mean residual life plot, showing a relatively linear plot up to the 0.95 threshold, indicating that this could be a good threshold. C. and D. show the parameter threshold stability plot of the modified scale and shape parameters. The dots demonstrate the scale and shape parameters for the Generalized Pareto distribution (GPD) at different thresholds. The lines show the 95% confidence interval. An optimal threshold is chosen at the point where both parameters are relatively stable, in this case, around the 95th percentile.*

### Extreme value theory results

In Figure 6, we show the data from two IDPs in the original space and after the marginal transformations have been applied. Furthermore, we show the histogram of $r$ exceeding the 0.95 quantile threshold. It can be noted that the variables fully lie on their respective axis after the marginal transformation.

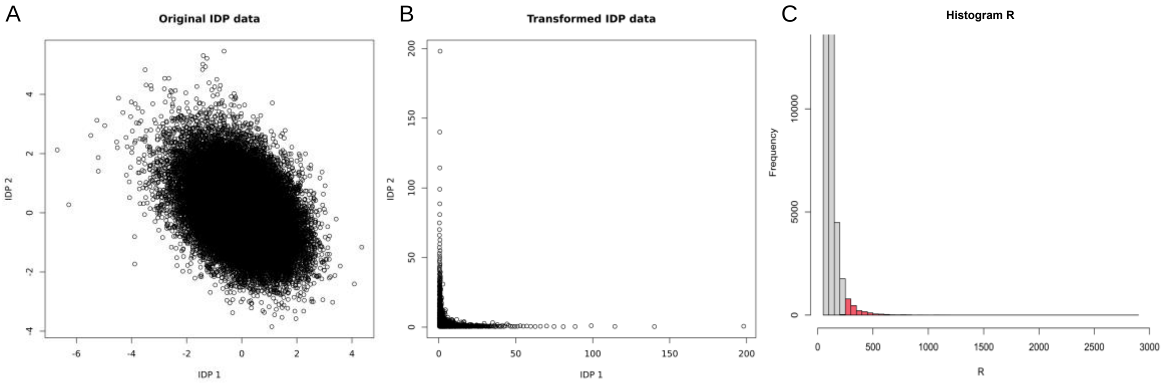


*Figure 6 - Illustration of the applied marginal transformation to the IDP data. A. Scatterplot of two IDPs in their original dimensions. B. Scatterplot of the IDP data after transforming the marginal, such that the marginal is regularly varying with α=2. C. Histogram showing the r polar coordinate and the 0.95 largest components.*

The $n_{exc} = 1960$ data is then used to estimate the TPDM $\widehat{\Sigma_{\mathbf{Z}}}$. In, Figure 7, we give an example of the found TPDM separated per IDP modality.

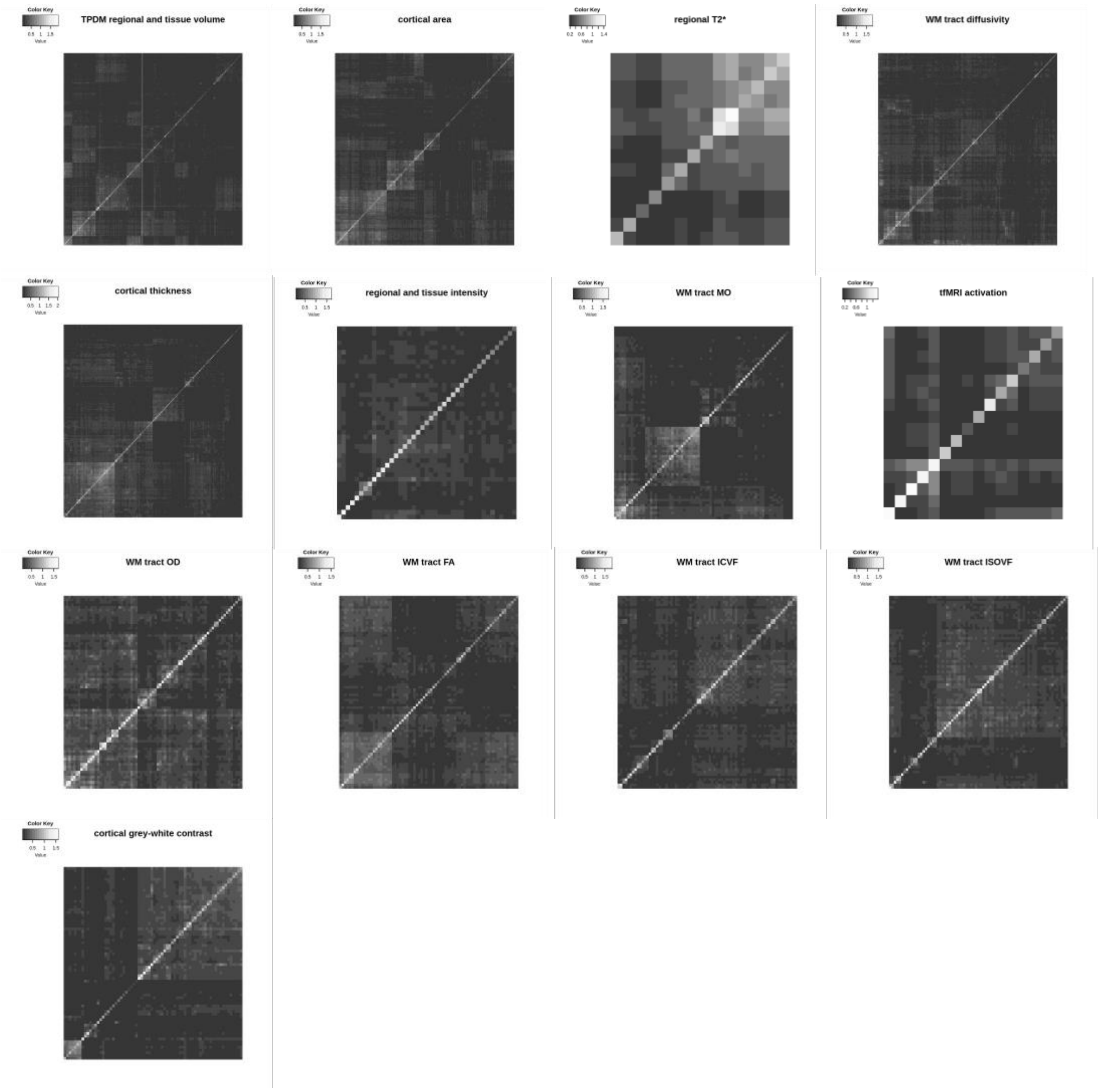


*Figure 7 - TPDMs separated per IDP modality.*

### Interpretation of extreme principal components

We performed a standard eigendecomposition of $\hat{\Sigma}_{\boldsymbol{z}}$ or the TPDM to obtain the eigenvectors (i.e. extreme PCs). Similar to covariance-PCA, we can look at the amount of scale that the first number of eigenvalues explains. In Figure 8, we show the correlation circle for the first two extreme PCs.

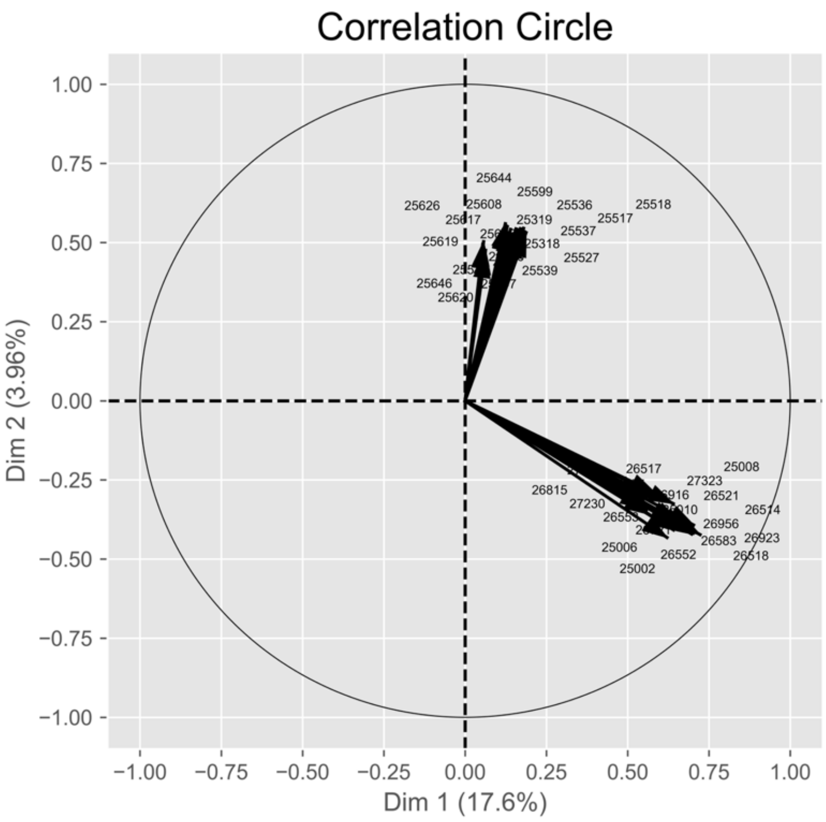


*Figure 8 – Showing the correlation circle with the loadings of the top 40 contributing variables on the extreme principal components 1 and 2.*

#### Removed IDPs

Since our objective was to demonstrate a new method, we removed a small number of IDPs which the likelihood warping approach did not fit well. An example is shown below.

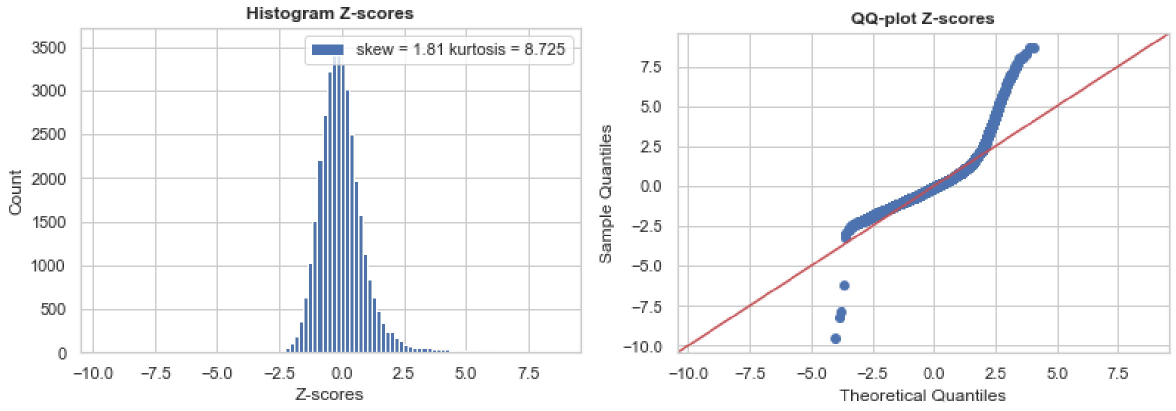


*Figure 9 - Showing an example of a badly fitted IDP with a high kurtosis. In the analysis, we have removed IDPs with a kurtosis above 10 from further processing, excluding one IDP.*

#### Standard PCs correlation with behavior

Here we show the results for correlating standard PCs to the behavioral variables. In Supplementary Figure 12, we present the Manhattan plots of the p-values for the univariate Spearman correlation between the nIDPs and the first two standard PCs. In Supplementary Table 1 we show a summary of the top nIDPs aggregated across both PCs.

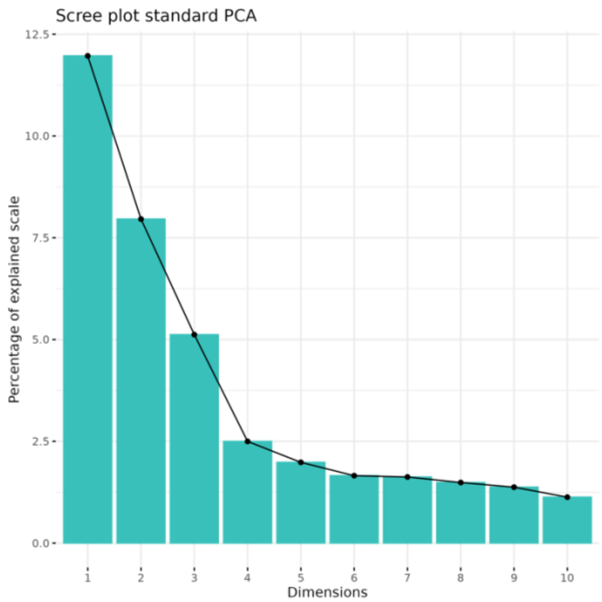


| | eigenvalue | scale.percent | cumulative.scale.percent |
|---|---|---|---|
| *Dim.1* | 245.96 | 11.97 | 11.97 |
| *Dim.2* | 163.58 | 7.96 | 19.93 |
| *Dim.3* | 105.24 | 5.12 | 25.05 |
| *Dim.4* | 51.35 | 2.5 | 27.55 |
| *Dim.5* | 40.76 | 1.98 | 29.53 |
| *Dim.6* | 34.04 | 1.66 | 31.19 |
| *Dim.7* | 33.38 | 1.62 | 32.81 |
| *Dim.8* | 30.57 | 1.49 | 34.3 |
| *Dim.9* | 28.24 | 1.37 | 35.67 |
| *Dim.10* | 23.2 | 1.13 | 36.8 |
| *Dim.11* | 20.92 | 1.02 | 37.82 |
| *Dim.12* | 20.68 | 1.01 | 38.83 |
| *Dim.13* | 20.18 | 0.98 | 39.81 |
| *Dim.14* | 18.63 | 0.91 | 40.72 |
| *Dim.15* | 17.37 | 0.85 | 41.56 |
| *Dim.16* | 16.43 | 0.8 | 42.36 |
| *Dim.17* | 15.01 | 0.73 | 43.09 |
| *Dim.18* | 14.81 | 0.72 | 43.81 |
| *Dim.19* | 13.68 | 0.67 | 44.48 |
| *Dim.20* | 13.02 | 0.63 | 45.11 |

*Figure 10 - Showing the scree plot of the standard PCA and table of scale values of the eigenvalue decomposition for the covariance matrix.*

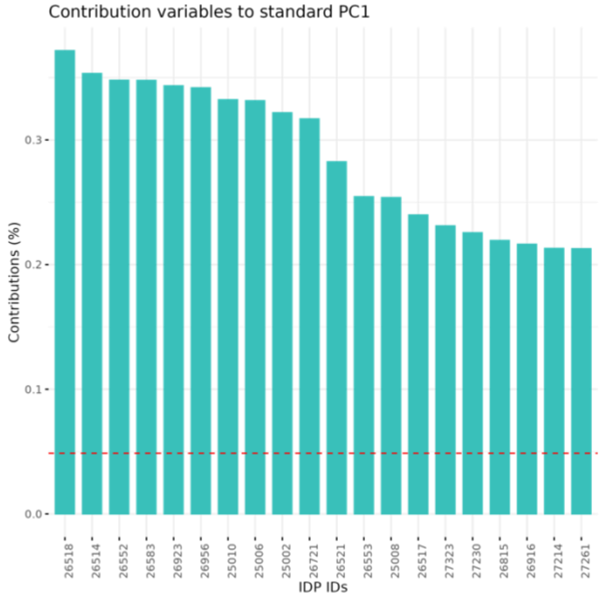


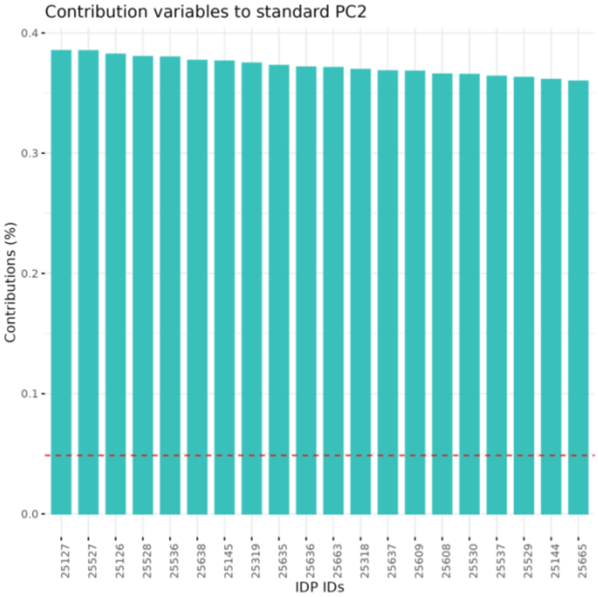


| Row.names | IDP.short.name | Category.name | contrib.1.top. |
|---|---|---|---|
| 26518 | aseg_global_volume_TotalGray | regional and tissue volume | 0.37 |
| 26514 | aseg_global_volume_BrainSeg | regional and tissue volume | 0.35 |
| 26552 | aseg_lh_volume_Cortex | regional and tissue volume | 0.35 |
| 26583 | aseg_rh_volume_Cortex | regional and tissue volume | 0.35 |
| 26923 | aparc-pial_lh_area_TotalSurface | cortical area | 0.34 |
| 26956 | aparc-pial_rh_area_TotalSurface | cortical area | 0.34 |
| 25010 | IDP_T1_SIENAX_brain-unnormalised_volume | regional and tissue volume | 0.33 |
| 25006 | IDP_T1_SIENAX_grey_unnormalised_volume | regional and tissue volume | 0.33 |
| 25002 | IDP_T1_SIENAX_peripheral_grey_unnormalised_volume | regional and tissue volume | 0.32 |
| 26721 | aparc-Desikan_lh_area_TotalSurface | cortical area | 0.32 |
| 26521 | aseg_global_volume_EstimatedTotalIntraCranial | regional and tissue volume | 0.28 |
| 26553 | aseg_lh_volume_CerebralWhiteMatter | regional and tissue volume | 0.25 |
| 25008 | IDP_T1_SIENAX_white_unnormalised_volume | regional and tissue volume | 0.25 |
| 26517 | aseg_global_volume_SubCortGray | regional and tissue volume | 0.24 |
| 27323 | aparc-DKTatlas_rh_volume_superiorfrontal | regional and tissue volume | 0.23 |
| 27230 | aparc-DKTatlas_lh_volume_superiorfrontal | regional and tissue volume | 0.23 |
| 26815 | aparc-Desikan_lh_volume_superiorfrontal | regional and tissue volume | 0.22 |
| 26916 | aparc-Desikan_rh_volume_superiorfrontal | regional and tissue volume | 0.22 |
| 27214 | aparc-DKTatlas_lh_volume_lateralorbitofrontal | regional and tissue volume | 0.21 |
| 27261 | aparc-DKTatlas_rh_area_superiorfrontal | cortical area | 0.21 |

| Row.names | IDP.short.name | Category.name | contrib.1.top. |
|---|---|---|---|
| 25127 | IDP_dMRI_TBSS_MD_Anterior_corona_radiata_L | WM tract diffusivity | 0.39 |
| 25527 | IDP_dMRI_ProbtrackX_MD_ifo_l | WM tract diffusivity | 0.39 |
| 25126 | IDP_dMRI_TBSS_MD_Anterior_corona_radiata_R | WM tract diffusivity | 0.38 |
| 25528 | IDP_dMRI_ProbtrackX_MD_ifo_r | WM tract diffusivity | 0.38 |
| 25536 | IDP_dMRI_ProbtrackX_MD_slf_l | WM tract diffusivity | 0.38 |
| 25638 | IDP_dMRI_ProbtrackX_L3_ilf_r | WM tract diffusivity | 0.38 |
| 25145 | IDP_dMRI_TBSS_MD_Superior_longitudinal_fasciculus_L | WM tract diffusivity | 0.38 |
| 25319 | IDP_dMRI_TBSS_L3_Anterior_corona_radiata_L | WM tract diffusivity | 0.37 |
| 25635 | IDP_dMRI_ProbtrackX_L3_ifo_l | WM tract diffusivity | 0.37 |
| 25636 | IDP_dMRI_ProbtrackX_L3_ifo_r | WM tract diffusivity | 0.37 |
| 25663 | IDP_dMRI_ProbtrackX_ICVF_ifo_r | WM tract ICVF | 0.37 |
| 25318 | IDP_dMRI_TBSS_L3_Anterior_corona_radiata_R | WM tract diffusivity | 0.37 |
| 25637 | IDP_dMRI_ProbtrackX_L3_ilf_l | WM tract diffusivity | 0.37 |
| 25609 | IDP_dMRI_ProbtrackX_L2_ifo_r | WM tract diffusivity | 0.37 |
| 25608 | IDP_dMRI_ProbtrackX_L2_ifo_l | WM tract diffusivity | 0.37 |
| 25530 | IDP_dMRI_ProbtrackX_MD_ilf_r | WM tract diffusivity | 0.37 |
| 25537 | IDP_dMRI_ProbtrackX_MD_slf_r | WM tract diffusivity | 0.36 |
| 25529 | IDP_dMRI_ProbtrackX_MD_ilf_l | WM tract diffusivity | 0.36 |
| 25144 | IDP_dMRI_TBSS_MD_Superior_longitudinal_fasciculus_R | WM tract diffusivity | 0.36 |
| 25665 | IDP_dMRI_ProbtrackX_ICVF_ilf_r | WM tract ICVF | 0.36 |

*Figure 11 - List of the top 20 contributions of the different IDPs to the first and second standard PCs.*

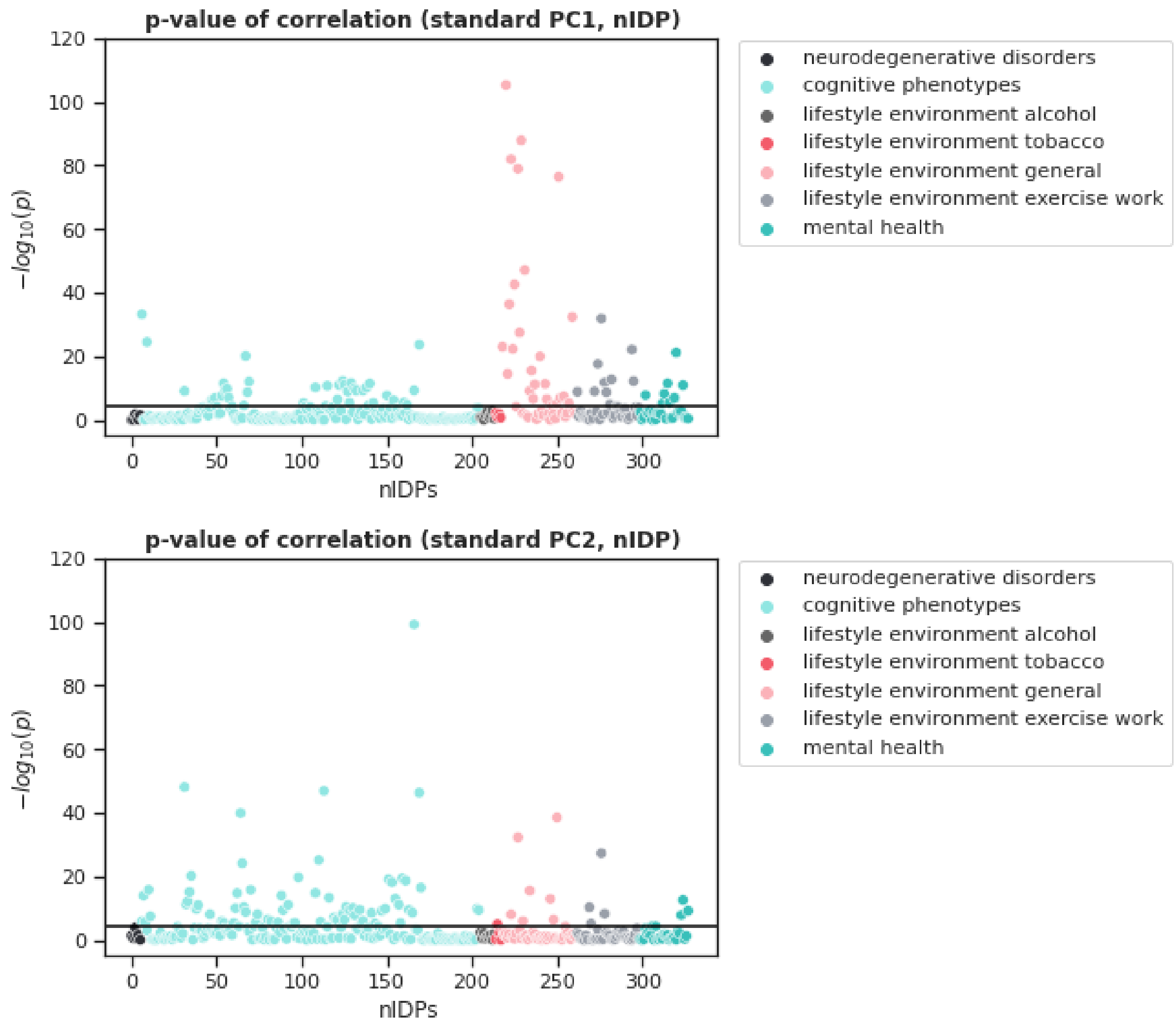


*Figure 12 - Showing two Manhattan plots of the log p-values for the Spearman correlation between the nIDPs and the first two standard PCs. The black line demonstrates the Bonferroni-corrected p-value threshold. Thus, values passing this line indicate a significant PC-nIDP correlation.*

| UKB nIDP | $-log_{10}(p)$ | Description |
|---|---|---|
| ***Neurodegenerative disorders*** | | |
| *No hits* | | |
| **Cognition** | | |
| 20016-2.0 | 99.2 | Fluid intelligence score |
| 630-2.0 | 48.1 | Touchscreen duration |
| 6373-2.0 | 46.9 | Number of puzzles correctly solved |
| 20128-2.0 | 46.4 | Number of fluid intelligence questions attempted within time limit |
| 4282-2.0 | 40.0 | Maximum digits remembered correctly |
| 398-2.3 | 33.2 | Number of correct matches in round |
| 6350-2.0 | 25.2 | Duration to complete alphanumeric path (trail #2) |
| 399-2.3 | 24.5 | Number of incorrect matches in round |
| 4283-2.0 | 24.2 | Number of rounds of numeric memory test performed |
| 4250-2.6 | 20.2 | Number of digits to be memorised/recalled |
| ***Lifestyle/environment alcohol*** | | |
| *No hits* | | |
| ***Lifestyle/environment tobacco*** | | |
| 20116-2.0 | 5.0 | Smoking status |
| ***Lifestyle/environment general*** | | |
| 20075-2.0 | 256.5 | Home location at assessment - north co-ordinate (rounded) |
| 4-2.0 | 166.4 | Biometrics duration |
| 5-2.0 | 105.3 | Sample collection duration |
| 1797-2.0 | 87.9 | Father still alive |
| 680-2.0 | 82.0 | Own or rent accommodation lived in |
| 738-2.0 | 79.0 | Average total household income before tax |
| 6142-2.0 | 76.5 | Current employment status |
| 1835-2.0 | 47.1 | Mother still alive |
| 709-2.0 | 42.6 | Number in household |
| 6138-2.0 | 38.5 | Qualifications |
| ***Lifestyle/environment exercise/work*** | | |
| 1070-2.0 | 31.9 | Time spent watching television (TV) |
| 6162-2.0 | 22.2 | Types of transport used (excluding work) |
| 1050-2.0 | 17.6 | Time spend outdoors in summer |
| 1130-2.0 | 12.7 | Hands-free device/speakerphone use with mobile phone in last 3 month |
| 6162-2.1 | 12.2 | Types of transport used (excluding work) |
| 1090-2.0 | 11.8 | Time spent driving |
| 924-2.0 | 10.3 | Usual walking pace |
| 981-2.0 | 9.0 | Duration walking for pleasure |
| 1011-2.0 | 8.8 | Frequency of light DIY in last 4 weeks |
| ***Mental health*** | | |
| 4537-2.0 | 21.2 | Work/job satisfaction |
| 4581-2.0 | 12.6 | Financial situation satisfaction |
| 2080-2.0 | 11.5 | Frequency of tiredness / lethargy in last 2 weeks |
| 4653-2.0 | 9.2 | Ever highly irritable/argumentative for 2 days |
| 2060-2.0 | 8.2 | Frequency of unenthusiasm / disinterest in last 2 weeks |
| 4570-2.0 | 7.8 | Friendships satisfaction |
| 1950-2.0 | 7.8 | Sensitivity / hurt feelings |
| 4526-2.0 | 6.9 | Happiness |
| 2090-2.0 | 5.4 | Seen doctor (GP) for nerves, anxiety, tension or depression |
| 2050-2.0 | 5.1 | Frequency of depressed mood in last 2 weeks |

*Table 1: Top 10 associations between non-imaging derived phenotypes (nIDPs) and standard principal components, grouped per category. The p-values reported in the table are the maximum p-value across the first two component. Only associations surviving Bonferroni-correction across nIDPs and components are reported.*
